# Stable White Matter Structure in the First Three Years after Psychosis Onset

**DOI:** 10.1101/2024.08.29.610312

**Authors:** Peter C. Van Dyken, Kun Yang, Andreia V. Faria, Akira Sawa, Michael MacKinley, Ali R. Khan, Lena Palaniyappan

## Abstract

**Background:** White matter alterations observed using diffusion weighted imaging have become a hallmark of chronic schizophrenia, but it is unclear when these changes arise over the course of the disease. Nearly all studies thus far have been cross-sectional, so despite their large sample sizes, they cannot determine if changes accumulate as a degenerative process, or if patients with pre-existing white matter damage are predisposed to more chronic forms of schizophrenia.

**Methods:** We examined 160 scans comprising two years of annual follow-up data from n=42 controls and n=28 schizophrenia patients recruited in the first two years since their diagnosis, totalling two to three scans per subject. We additionally examined six-month follow-up data obtained from an ultra-high field (7 Tesla) scanner (n=68 scans; n=19 first-episode schizophrenia patients; n=15 controls) as a validation dataset. A longitudinal model was used to compare the trajectory of diffusion tenor parameters between patients and controls. Positive and negative symptom scores were correlated with diffusion parameters using ROI- and clustering-based approaches.

**Results:** We failed to observe any longitudinal differences in any diffusion tensor imaging parameters between patients and controls in either dataset. We did, however, observe consistent associations between white matter alterations and negative symptoms in both datasets.

**Conclusions:** White matter does not appear susceptible to schizophrenia-linked degeneration in the early stages of disease, but pre-existing pathology may be linked to disease severity.

## 1 Introduction

Studies of chronic patients using diffusion tensor imaging (DTI) have robustly demonstrated a reduction of fractional anisotropy (FA) across the white matter, potentially reflecting increased axonal disorganization or reduced fibre integrity. However, these findings are much more limited in patients with early psychosis. Our own recent investigation failed to find any substantial differences between patients and controls in two separate datasets (1). Other studies have found differences only with limited effect size (2–14). The neurodegenerative hypothesis of white matter in schizophrenia argues that patients experience progressive deterioration of white matter over the disease course. It is possible that despite the lack of substantial white matter integrity at the outset, DTI metrics may begin to change after the disease onset.

Previous reports have argued in favour of this white matter neurodegeneration hypothesis, citing data showing increasingly disrupted DTI metrics in more chronic patients (15). Nearly all of these studies, however, have been cross-sectional, and though they all report “declining” FA in chronic patients (16–18), they cannot disambiguate individual changes over time from cohort effects. In other words, rather than degenerating over time, patients with pre-existing lower FA may be selectively progressing to chronic forms of schizophrenia. Cross-sectional data are inadequate to test this alternative explanation.

Unfortunately, high quality longitudinal studies with diffusion imaging, including both patients and controls, followed for multiple years, are rare. Regarding chronic patients, Tronchin et al. (19) have reported higher decline in FA in patients after six months around the genu of the corpus callosum, whereas Domen et al. (20) failed to find any group by time differences in a three-year follow-up study of 55 patients, although they did note both relative increases and decreases of FA in patients compared to healthy controls (HCs) in a few ROIs. In first episode psychosis (FEP) patients, Berge et al. (21) failed to find an effect of time in 20 patients after a two-year follow-up; Serpa et al. (10) reached a similar conclusion in a six-month follow-up of 21 patients. Results from ultra-high-risk patients (without a formal schizophrenia diagnosis) have been heterogeneous, with two studies finding significant decreases in FA compared to controls (22,23), one finding a significant increase over six months (24), and three failing to find any differences (25–27). All other studies reviewed were limited by lack of HC follow-up, a follow-up time of less than six months, or a lack of angular resolution in the diffusion sequence (less than 10 directions) precluding reliable inference.

In HCs, FA peaks in most regions by around age thirty, then gradually declines with age (28–31). Previous authors have suggested this age-related decline starts early in schizophrenia patients (15,32), resulting in consistently lower FA across their adult life, even though the absolute rate of decline may be similar to HCs. Thus, the early stages of psychosis are likely the most important for detecting individual effects on FA trajectory. With only two, relatively small, published studies investigating such patients, we cannot definitively say whether this early peak exists. In this report, we address this gap with findings from the Johns Hopkins Schizophrenia Center (JHSZC) dataset, a two-year follow-up study of early psychosis patients and HCs with three time points for most participants. We also validate our analysis with six-month follow-up data from the Tracking Outcomes in Psychosis (TOPSY) dataset previously reported (1). Given the age of our participants is close to the early-peak period of developmental risk (15), we expected to find significant FA decline (with mean diffusivity (MD), radial diffusivity (RD), and axial diffusivity (AD) increases (33)) compared to HCs.

## 2 Materials and Methods

### 2.1 Data

#### 2.1.1 JHSZC

This study was approved by the Johns Hopkins Medicine Institutional Review Board. All study participants provided written informed consent. 101 patients and 96 controls were recruited from within Johns Hopkins Hospital and the surrounding region. Inclusion criteria were: 1) between 13 and 35 years old; 2) no history of traumatic brain injury, cancer, abnormal bleeding, viral infection, neurologic disorder, or intellectual disability; 3) no drug or alcohol abuse (not including cannabis or synthetic cannabinoid receptor agonists) in the past three years; 4) no illicit drug use in the past two months. Patients were within 24 months of the onset of psychotic symptoms as assessed by study team psychiatrists. Subjects diagnosed with bipolar disorder with psychotic features (n=22), major depressive disorder with psychotic features (n=4), substance induced psychotic disorder (n=3), and psychosis not otherwise specified (n=3) were excluded.

Follow-up assessments and imaging for both HCs and patients were obtained where possible at one, two, and three years from baseline.

Data was acquired with 3T MRI (Philips Intera (v3.2.2)) using T1-weighted (T1w) and diffusion magnetic resonance imaging (dMRI) imaging protocols. T1w data was collected using an MPRAGE sequence at 1 mm isotropic resolution. Two replicate diffusion datasets were acquired per subject per session with an echo planar imaging (EPI) sequence at 0.74x0.74x2.2 mm resolution. 32 directions were acquired in the AP direction at b=700, along with 1 b=0 images (details in supplementary methods).

Baseline, but not follow-up, diffusion data from this dataset has been previously reported (34–36).

#### 2.1.2 TOPSY

FEP patients and HCs were recruited from an established cohort enrolled in the Prevention and Early Intervention Program for Psychoses (PEPP) in London, Ontario. Inclusion criteria for FEP patients was as follows: individuals experiencing their first psychotic episode, with no more than 14 days of cumulative lifetime antipsychotic exposure, no major head injuries, no known neurological disorders, and no concurrent substance use disorder. All participants provided written, informed consent prior to participation. Patient diagnosis was established using a best estimate procedure (37) and confirmed after 6 months of treatment. Two patients with major depressive disorder and two with bipolar disorder were excluded from analysis.

Follow-up assessments and imaging for both HCs and patients were obtained where possible at six months, one year, two years, and three years.

Data was acquired with a head-only, neuro-optimized 7T MRI (Siemens MAGNETOM Plus, Erlangen, Germany) using T1w and dMRI imaging protocols. T1w data was collected using an MP2RAGE sequence (38) at 0.75 mm isotropic resolution. Diffusion data was acquired with an EPI sequence at 2mm isotropic resolution. 64 directions were acquired in both the AP and PA directions at b=1000, along with 2 b=0 images (details in supplementary methods). Gradient nonlinearity correction was applied to all acquisitions using in-house software.

Baseline, but not follow-up, diffusion data from this dataset has been previously reported (1).

### 2.2 Preprocessing

Anatomical and diffusion preprocessing for TOPSY data was performed as previously reported (1). Diffusion processing included denoising; susceptibility distortion correction with reverse phase-encoded images; motion correction; registration to the T1 image; fitting DTI tensors; probabilistic tractography using constrained spherical deconvolution; and generation of cortical connectomes using the Brainnetome atlas (39), weighted using sampled DTI parameters.

JHSZC data were preprocessed as above with the following deviations:

The replicate diffusion acquisitions were concatenated before processing. Because we did not have a reverse phase-encoding scan, susceptibility distortion correction was performed using an approach mediated by *SynthSR* (40) and *antsRegistration* (41), as described in the supplementary methods. Slice to volume correction in *eddy* was used to correct intravolume motion correction. Volumes with severe artefacts were manually excluded. The DTI tensor was fit in the original acquisition space.

Full preprocessing details are in the supplementary methods.

### 2.3 Parcellations

FA maps were non-linearly registered to a common template corresponding to the average space of all FA images.

FA, MD, AD, and RD values were projected to an FA-derived skeleton, then sampled with a composite atlas comprising the Johns Hopkins University (JHU) white matter atlas, capturing core white matter regions, and the Talairach lobe segmentation, capturing peripheral white matter regions. To increase our sensitivity to local and global changes, the ROIs were organized into a hierarchical system. The first level was a single ROI covering the entire white matter skeleton. The second split the white matter into core and peripheral regions. The third included the peripheral lobe ROIs derived from the Talairach segmentation and four groupings of the JHU atlas labelled as projection, association, callosal, and limbic tracts (see Table 2). The final level included the individual JHU ROIs. Multiple comparisons were corrected separately across each level using the Benjamini-Hochberg false discovery rate (FDR) procedure (42). This approach was based on previous work by Simmonds et al. (43).

**Table 1:**
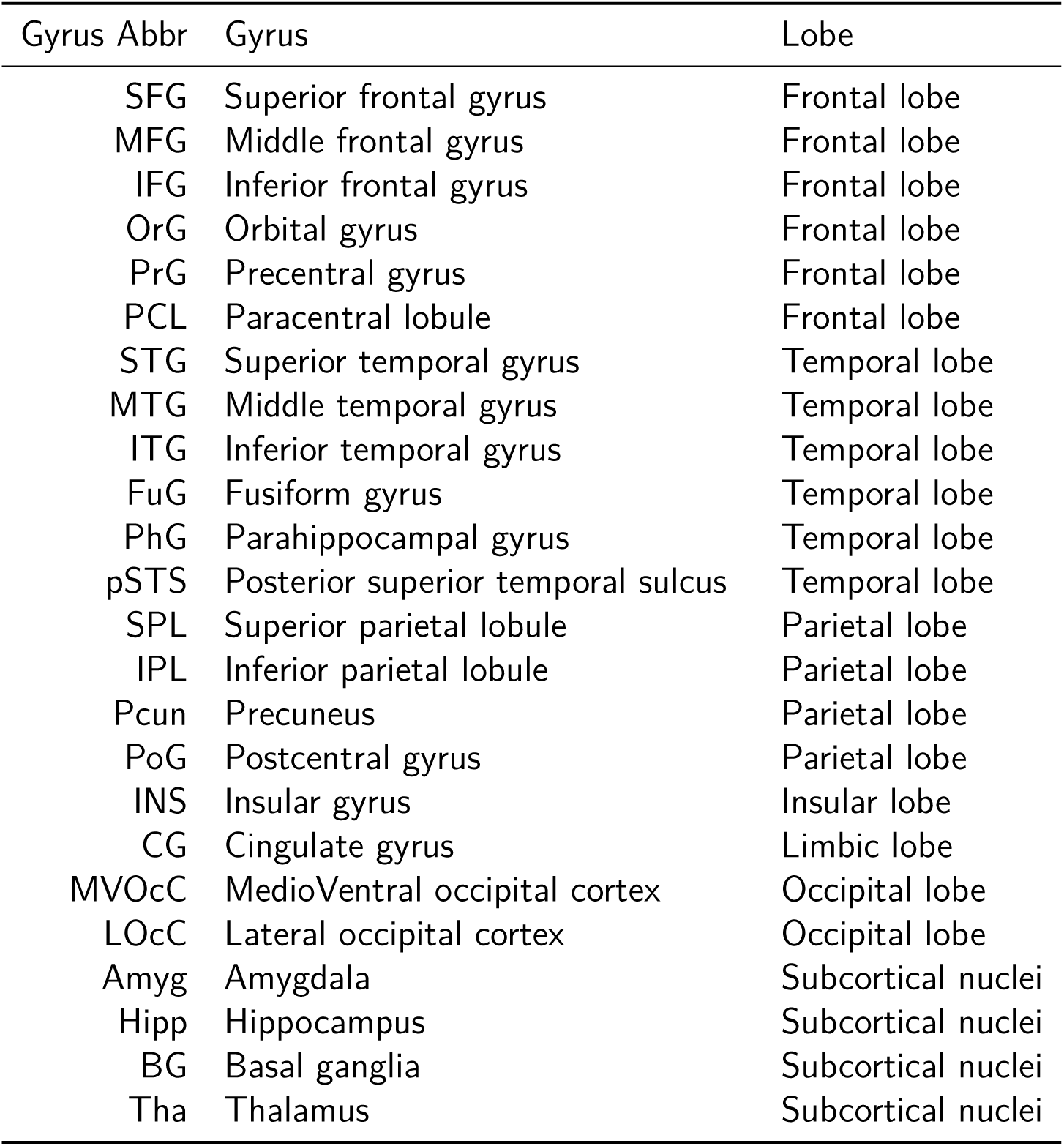
Abbreviations of cortical regions.

| Gyrus Abbr | Gyrus | Lobe |
| --- | --- | --- |
| SFG | Superior frontal gyrus | Frontal lobe |
| MFG | Middle frontal gyrus | Frontal lobe |
| IFG | Inferior frontal gyrus | Frontal lobe |
| OrG | Orbital gyrus | Frontal lobe |
| PrG | Precentral gyrus | Frontal lobe |
| PCL | Paracentral lobule | Frontal lobe |
| STG | Superior temporal gyrus | Temporal lobe |
| MTG | Middle temporal gyrus | Temporal lobe |
| ITG | Inferior temporal gyrus | Temporal lobe |
| FuG | Fusiform gyrus | Temporal lobe |
| PhG | Parahippocampal gyrus | Temporal lobe |
| pSTS | Posterior superior temporal sulcus | Temporal lobe |
| SPL | Superior parietal lobule | Parietal lobe |
| IPL | Inferior parietal lobule | Parietal lobe |
| Pcun | Precuneus | Parietal lobe |
| PoG | Postcentral gyrus | Parietal lobe |
| INS | Insular gyrus | Insular lobe |
| CG | Cingulate gyrus | Limbic lobe |
| MVOcC | MedioVentral occipital cortex | Occipital lobe |
| LOcC | Lateral occipital cortex | Occipital lobe |
| Amyg | Amygdala | Subcortical nuclei |
| Hipp | Hippocampus | Subcortical nuclei |
| BG | Basal ganglia | Subcortical nuclei |
| Tha | Thalamus | Subcortical nuclei |

**Table 2:** ROIs used in study, with the four subgroupings of the JHU atlas.

| Atlas | Group | Region |
| --- | --- | --- |
| Core (JHU) | Association | External capsule |
|  |  | Inferior fronto-occipital fasciculus |
|  |  | Sagittal stratum |
|  |  | Superior fronto-occipital fasciculus |
|  |  | Superior longitudinal fasciculus |
|  |  | Uncinate fasciculus |
|  | Callosal | CC Body |
|  |  | CC Genu |
|  |  | CC Splenium |
|  | Limbic | Cingulate gyrus |
|  |  | Fornix |
|  |  | Fornix (cres) |
|  |  | Hippocampus |
|  | Projection | Anterior corona radiata |
|  |  | Cerebral peduncle |
|  |  | Corticospinal tract |
|  |  | IC Anterior Limb |
|  |  | IC Posterior Limb |
|  |  | IC Retrolenticular |
|  |  | Medial lemniscus |
|  |  | Posterior corona radiata |
|  |  | Posterior thalamic radiation |
|  |  | Superior corona radiata |
| Peripheral |  | Anterior Lobe |
|  |  | Frontal Lobe |
|  |  | Limbic Lobe |
|  |  | Occipital Lobe |
|  |  | Parietal Lobe |
|  |  | Posterior Lobe |
|  |  | Temporal Lobe |

TOPSY, we additionally split the subjects into two groups according to whether their score was at the minimum value ( $=3$ ) at the second session (this was done separately for each score). The minimal-score group was called the “remission” group. ROI statistics were performed using T-tests, controlling for age and sex. One-way comparisons were used, testing that lower FA, and higher MD, AD, and RD, correlated with higher score slope and intercept. For remission, we tested that FA was higher, and MD, AD, and RD lower, in the recovery group.

### 2.4 Analysis

Longitudinal analysis was performed with a linear mixed effects model, implemented in R [v4.3.3; (44)] using *lme4* [v1.1.35.3; (45)]. Age and sex were regressed as nuisance variables, and random intercepts and slopes were fit for each subject (only random slopes for TOPSY dataset, which had just two timepoints). The full model was as follows:

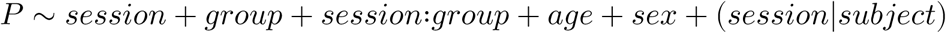

Significance of the random slopes and intercepts was tested using *ranova* from *lmerTest* [v3.1.3; (46)]. Degrees of freedom for the main each comparison was estimated using Satterthwaite’s method (47) with *lmertest* [v3.1.3; (46)]. For FA, a one-way comparison was performed testing for more pronounced FA reductions in patients than controls. For MD, AD, and RD, a one-way comparison was performed in the opposite direction.

For clinical score analyses, the sampled ROIs, parameter images, and connectomes were averaged across sessions for each subject. Positive and negative clinical scores were considered. For score and subject, the intercept and slope were computed by fitting the scores across sessions to a first-order polynomial. For TOPSY, we additionally split the subjects into two groups according to whether their score was at the minimum value (=3) at the second session (this was done separately for each score). The minimal-score group was called the “remission” group. ROI statistics were performed using T-tests, controlling for age and sex. One-way comparisons were used, testing that lower FA, and higher MD, AD, and RD, correlated with higher score slope and intercept. For remission, we tested that FA was higher, and MD, AD, and RD lower, in the recovery group.

Tract-Based Spatial Statistics (TBSS) was performed using the skeletonized images and *FSL randomise* (48) with threshold-free cluster enhancement-correction and 10,000 permutations, thresholding at a corrected p-value of 0.05. Sex and age were regressed as nuisance variables. One-way comparisons were performed as in the ROIs.

Network based statistic (NBS) was performed with extent-based cluster sizes, a T threshold of 3.0, 10,000 iterations, and corrected p-value threshold of 0.05. Sex and age were regressed as nuisance variables (49). One-way comparisons were performed as in the ROIs.

### 2.5 Code Availability

Except where otherwise indicated, all analyses were performed using Python 3.11. The analyses discussed above and resulting figures were made possible by openly available python packages (50–65). All code used is freely available at https://github.com/pvandyken/study-jhp_topsy_longt.git. Links to pipelines used for data preprocessing are listed at that repository.

## 3 Results

### 3.1 Demographics

In the JHSZC dataset (Table 3), 50 HCs and 40 patients were lost to follow-up (Table S3). Four additional HCs and one patient did not have usable diffusion data. This left 42 HCs and 24 patients from the JHSZC dataset. Additionally, two session-specific scans were dropped due to quality, but the affected subjects still had two usable sessions included in the study. The number of dropouts per group did not vary (𝜒^2^(1) = 1.17; 𝑃 = .28).

**Table 3:** JHSZC demographics.

|  | Healthy Control (n=42) |  |  | Early Psychosis (n=28) |  |  |  |
| --- | --- | --- | --- | --- | --- | --- | --- |
|  | Baseline<br>(n=42) | 1yr<br>(n=37) | 2yr<br>(n=18) | Baseline<br>(n=24) | 1yr<br>(n=25) | 2yr<br>(n=13) | 3yr<br>(n=1) |
| Sex (M/F) | 22/20 | 20/17 | 9/9 | 19/5 | 20/5 | **12/1** | 1/0 |
| Age | 23.93<br>(3.39) | 25.30<br>(3.39) | 25.72<br>(3.48) | **21.88**<br>**(4.00)** | **23.00**<br>**(3.85)** | **22.15**<br>**(2.91)** | 24.00 |
| Ethnicity<br>(B/EA/O/W) | 23/2/2/15 | 20/1/2/14 | 10/1/1/6 | 17/0/1/6 | 19/0/1/5 | 10/0/0/3 | 1/0/0/0 |
| Handedness<br>(R/L) | 35/7 | 30/7 | 17/1 | 21/3 | 22/3 | 11/2 | 1/0 |
| Smoker<br>(Yes/No) | 3/39 | 4/33 | 1/17 | 5/19 | **12/13** | 5/8 | 0/1 |
| Cannabis<br>(Yes/No) | 3/39 | 4/32 | 2/16 | 4/20 | 6/19 | 0/13 | 0/1 |
| Duration of<br>Illness (weeks) |  |  |  | 65.00<br>(46.58)<sup>†</sup> | 121.33<br>(56.33)<sup>†</sup> | 160.33<br>(26.00)<sup>†</sup> | 255.67<sup>†</sup> |
| CPZ (mg) |  |  |  | 253.33<br>(234.29) | 282.76<br>(277.64) | 359.62<br>(332.84) | 250.00 |
| SAPS |  |  |  | 4.48<br>(4.36) | 4.48<br>(2.29) | 3.62<br>(4.39) | 0.00 |
| SANS |  |  |  | 8.43<br>(4.04) | 7.61<br>(3.50) | 7.31<br>(5.71) | 0.00 |

<sup>†</sup> Median (IQR)
B=Black/African; EA=East Asian; O=Other/Unknown; W=White/European; CPZ=chlorpromazine equivalent dose; SAPS=Scale for assessment of positive symptoms; SANS=Score for assessment of negative symptoms

For the included subjects, the distribution of HCs and patients across sessions did not significantly vary (𝜒^2^(2) = 0.41; 𝑃 = .81). There was a significantly greater proportion of males than females at each session in the patient group than the control group, and controls were significantly older than patients across the study. Patients who returned for their one-year follow-up were also significantly more likely to smoke than their healthy counterparts (Table S1). Initial follow-ups were 1.1 ± 0.1 years following the baseline scan. Second follow-ups were 2.1 ± 0.2 years following the baseline scan. No significant difference in follow-up time was observed between patients and HCs for the first (𝑇 (62) = 0.29; 𝑃 = .77) or second (𝑇 (30) = 0.93; 𝑃 = .36) follow-up (Figure S1 A).

In the TOPSY dataset (Table 4), 23 HCs and 49 patients were lost to follow-up (Table S4). One HC and three patients did not have usable diffusion data from each session, leaving 15 HCs and 19 patients for analysis. The number of dropouts did not significantly vary between groups (𝜒^2^(1) = 0.34; 𝑃 = 0.56). Second follow-ups were obtained for four patients, but not for any controls, thus these data were excluded from the study.

**Table 4:** TOPSY demographics.

|  | Healthy Control |  | First Episode Psychosis |  |
| --- | --- | --- | --- | --- |
|  | Baseline (n=15) | 6 mo (n=15) | Baseline (n=19) | 6 mo (n=19) |
| Sex (M/F) | 10/5 | 10/5 | 16/3 | 16/3 |
| Age | 21.73 (2.99) | 22.60 (3.04) | 22.53 (5.42) | 23.11 (5.40) |
| Handedness<br>(R/L/A) | 14/0/1 | 14/0/1 | 18/0/1 | 18/0/1 |
| Education | 14.27 (2.02) | 14.33 (2.09) | **12.95 (1.18)** | **12.95 (1.18)** |
| SES | 3.20 (1.57) | 3.20 (1.57) | 3.89 (1.08) | 3.89 (1.08) |
| CAST | 7.00 (3.87) | 13.50 (10.61) | **12.12 (6.90)** |  |
| AUDIT-C | 2.87 (2.26) | 5.00 (2.83) | 3.79 (3.83) |  |
| Smoker<br>(yes/no) | 1/14 | 1/14 | 7/12 | 0/19 |
| Cannabis<br>(yes/no) | 5/10 | 1/1 | 12/7 | 0/19 |
| SOFAS | 81.08 (6.24) | 79.71 (5.25) | **42.00 (13.37)** | **61.89 (14.68)** |
| Duration of<br>Illness (weeks) |  |  | 104.00 (132.50)<sup>†</sup> | 130.07 (140.18)<sup>†</sup> |
|  | Baseline (n=15) | 6 mo (n=15) | Baseline (n=19) | 6 mo (n=19) |
| Antipsych.<br>Day of Scan<br>(DDD) |  |  | 0.00 (0.33)<sup>†</sup> | 1.00 (0.66)<sup>†</sup> |
| Antipsych.<br>Lifetime<br>(DDD-days) |  |  | 0.00 (1.25)<sup>†</sup> | 135.64 (107.44)<sup>†</sup> |
| CDS |  |  | 0.58 (0.51) | 0.21 (0.42) |
| PANSS-8<br>Total |  |  | 24.68 (5.53) | 13.53 (4.57) |
| PANSS-8<br>Positive |  |  | 12.53 (2.82) | 4.95 (1.93) |
| PANSS-8<br>Negative |  |  | 6.26 (3.31) | 5.95 (3.58) |
| PANSS-8<br>General |  |  | 5.89 (2.54) | 2.53 (1.12) |

<sup>†</sup> Median (IQR)
B=Black/African; C=Caribbean/North American Black; EA=East Asian; W=White/European; DDD=Defined daily dose; DDD-days=DDD × Days; CDS=Calgary Depression Scale; CAST=Cannabis Abuse Screening Test; PANSS=Positive and Negative Symptom Scale; SES=Socioeconomic status; AUDIT-C=Alcohol Use Disorders Identification Test; SOFAS=Social and Occupational Functioning Assessment Scale

HCs and patients were matched for age, sex, and socioeconomic status. Patients had significantly lower levels of education, had a higher Cannabis Abuse Screening Test (CAST) score indicating increased risk of cannabis abuse (66), and had a significantly lower Social and Occupational Functioning Assessment Scale (SOFAS) score (67) (Table S2). There was no correlation between the Calgary Depression Score and the PANSS8-N subscore. Follow-ups were 0.6 ± 0.3 years following the baseline scan. HCs had significantly longer follow-up times than patients (𝑇 (35) = 2.44; 𝑃 = .020) (Figure S1 B).

### 3.2 DTI metrics remain stable across time

To investigate longitudinal changes of DTI parameters in first episode patients, FA, MD, AD, and RD values were sampled from a comprehensive white matter atlas, constructed as described in Section 2.3. A mixed linear effects paradigm was used to model parameter changes across sessions. Subjects were used as a random effect and age, sex, group, and session were used as fixed effects.

No significant differences in slope between healthy controls and patients were observed for any parameter in any ROI, after correcting for multiple comparisons (Figure 1). A significant increase in global FA (two-tailed test; 𝑇 (132.50) = 2.06; 𝑃 = .042) and decrease in RD (𝑇 (130.00) = −2.01; 𝑃 = .047) was observed across all subjects in the JHU dataset.

**Figure 1:**
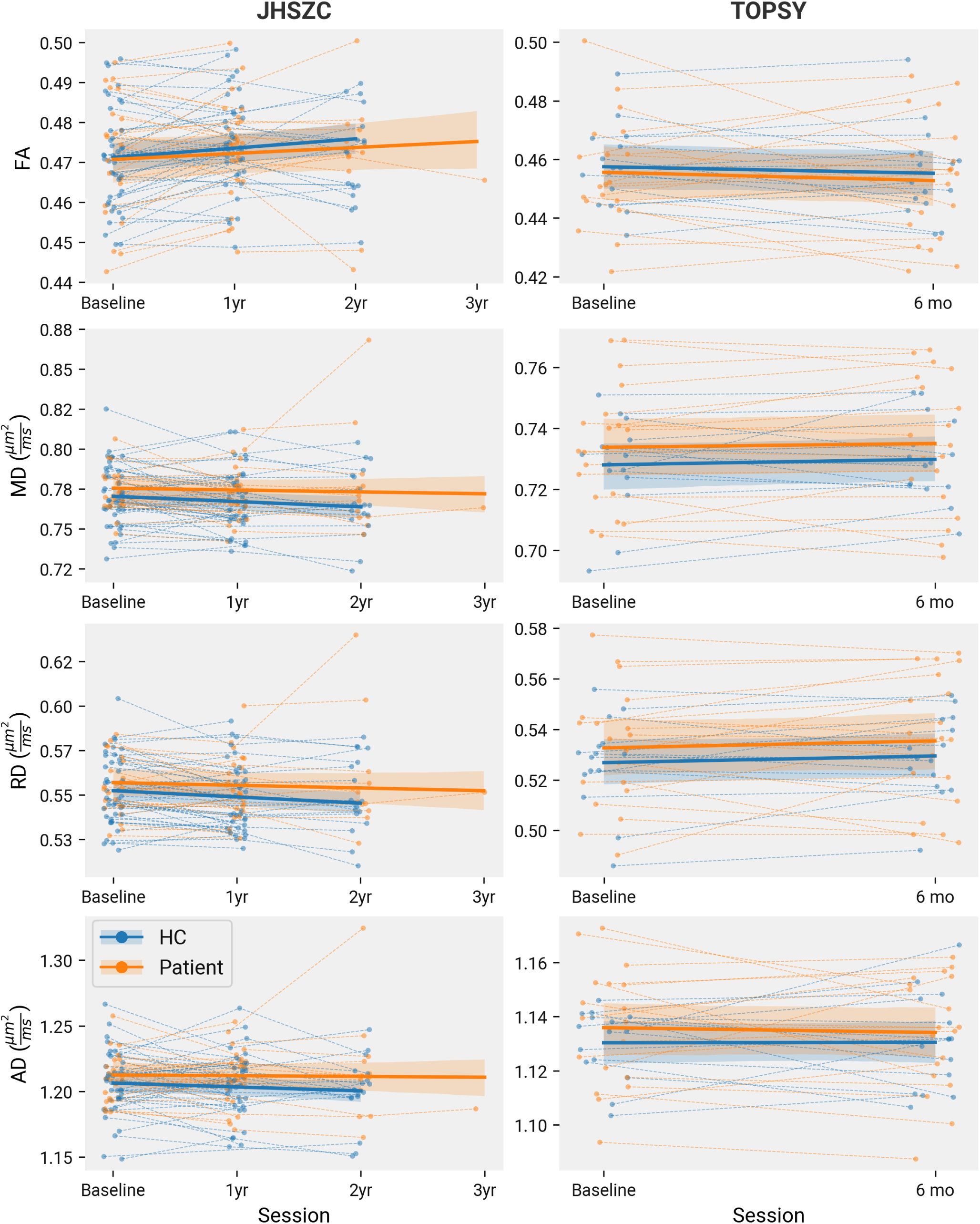
Global longitudinal changes of white matter microstructure in early schizophrenia patients. Trendlines show a linear mixed effect model of parameter against session with random intercepts fit for every subject. Shaded bands show a 95% CI computed with parametric bootstrapping resampling residuals and random effects 1000 times. No significant differences were found between the slopes of HCs and patients in either dataset for any of the parameters measured. In the JHSZC sample, fitting random slopes to each subject did not significantly improve the fit of the model (not tested in TOPSY because each subject had only two time points).

### 3.3 Age-matched sub-study in JHSZC

To explore if the lack of findings in the JHSZC dataset could be explained by the differing age and sex distributions between the HCs and patients, we used a single age cut-off across the dataset, selecting subjects younger than 24 years. The study groups in the resulting dataset were matched for age and sex (Table S5, Table S6). Repeating the above mixed linear effects model, no differences in longitudinal change were found between groups. The global increase and decrease of FA and RD were also no longer observed (Figure 2).

**Figure 2:**
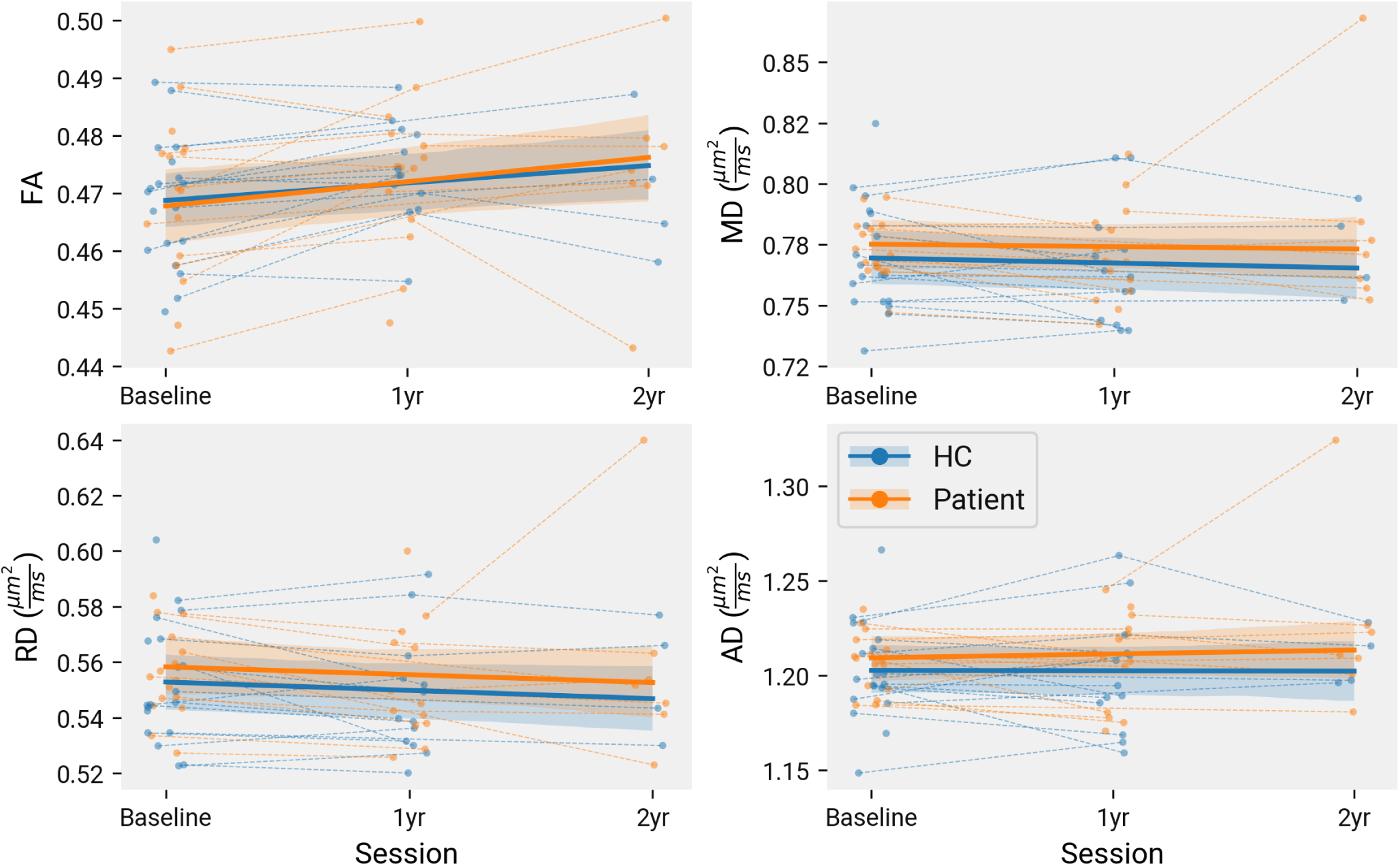
Global longitudinal changes of white matter microstructure in age-matched subset of JHSZC dataset. Trendlines show a linear mixed effect model of parameter against session with random intercepts fit for every subject. Shaded bands show a 95% CI computed with parametric bootstrapping resampling residuals and random effects 1000 times. No significant differences were found between the slopes of HCs and patients for any of the parameters measured. Fitting random slopes to each subject did not significantly improve the fit of the model.

### 3.4 DTI metrics in JHSZC

We next investigated whether the clinical scores corresponded with DTI metrics. Positive and negative symptoms were measured in each dataset, in JHSZC using Scale for the Assessment of Positive Symptoms (SAPS) and Scale for the Assessment of Negative Symptoms (SANS) and in TOPSY using PANSS8-P and PANSS8-N. For these analyses, we included all patients with valid clinical score measurements from at least two sessions and at least one valid scan. This gave us n=28 for the JHSZC dataset (identical to the longitudinal analysis) and n=21 for TOPSY (two additional subjects with valid clinical scores but missing scans at their second session). To characterize the change and baseline of clinical scores, we fit a first-order linear model to the positive and negative scores of each subject and extracted the slopes and intercepts. We then modelled these against the average DTI metrics across our ROIs, using age and sex as covariates. Because we did not find significant subject-specific slopes in our longitudinal analysis, we averaged the DTI data from each subject across available sessions. As in previous reports, we expected worse symptom baselines and trajectories to correspond with lower FA and higher MD, RD, and AD. We therefore performed inference using 1-tailed t-tests in the corresponding directions for all following tests.

In the JHSZC dataset, comprising patients enrolled after treatment commencement, neither clinical score had significant change across session (Figure 3 A). The SANS intercept significantly correlated with the global average of DTI metrics: higher symptom severity corresponded with lower FA and higher with MD, RD, and AD. The four significant interactions were reproduced in the core white matter ROI. The MD, RD, and AD interactions were observed in the peripheral white matter ROI (Figure 4). Interactions between MD and RD and the SANS intercept were further observed in the frontal, limbic, posterior, and temporal lobe ROIs and in the association, callosal, and projection ROI groups. AD interactions were found in frontal and temporal lobe and association tract ROIs (Figure 4). Finally, the same parametershad interactions with several of the JHU atlas ROIs, as tabulated in Table 5 and visualized in Figure 5. FA had no further significant interactions. None of the parameters had interactions with the SANS slope in any ROI, or any interactions with the SAPS slope or intercept.

**Figure 3:**
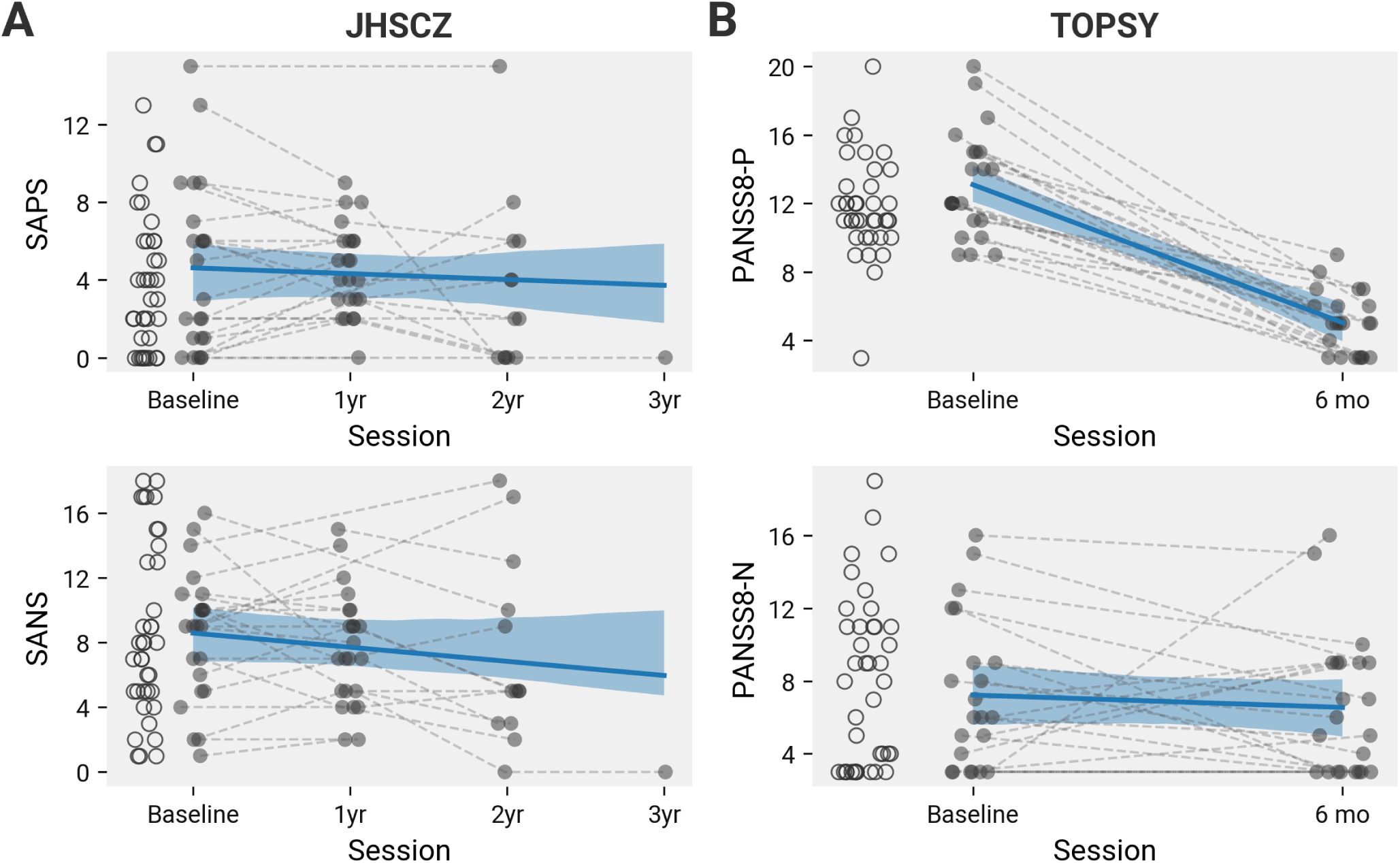
Baseline and follow-up clinical scores in early schizophrenia patients. At baseline, empty circles show subjects with no follow-ups. Filled circles connected by a line represent the same subject across multiple visits. No significant differences in baseline scores were found between subjects with and without follow-up visits. Trendlines show a linear fixed effect model of parameter against session with random slopes and intercepts fit for every subject (only random intercepts for TOPSY). Shaded bands show a 95% CI computed with parametric bootstrapping resampling residuals and random effects 500 times. In the TOPSY dataset, the PANSS8-P score was significantly lower in the second session than the first (1000 perms, p < .001). No other symptom scores significantly changed across session.

**Figure 4:**
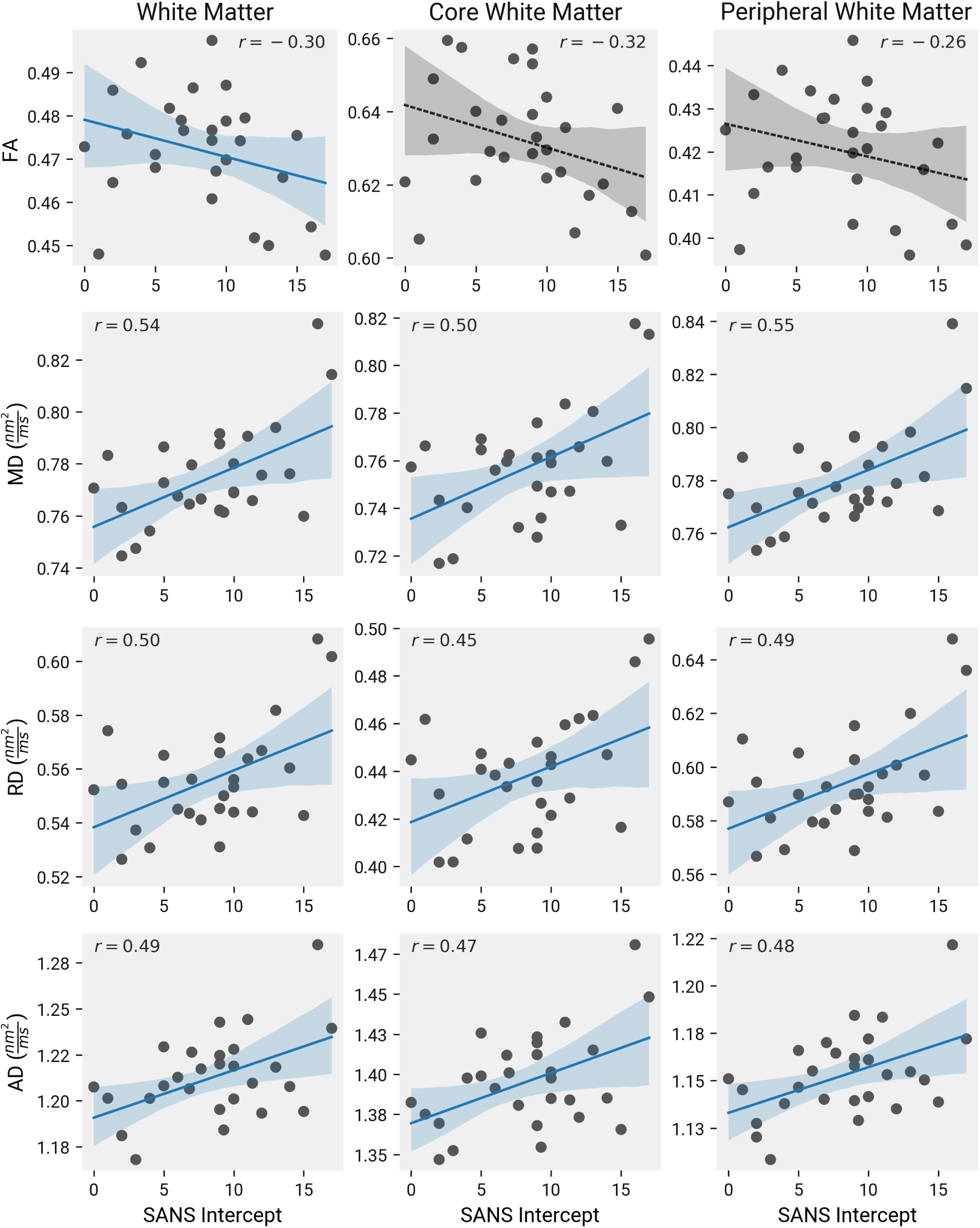
Correlation between microstructural parameters and negative symptom intercepts in the JHSZC dataset. Microstructural measurements are averaged across all sessions for each subject. The negative symptom (SANS) intercept was computed using a first order linear model for each subject, with the baseline scan as time 0. Shaded bands show 95% CI computed with nonparametric bootstrap paired resampling with 1000 permutations. Relationships were tested with a linear model with age and sex and covariates. Solid lines represent statistically significant relationships, dashed lines are nonsignificant. T-values and P-values are shown in Table 5.

**Figure 5:**
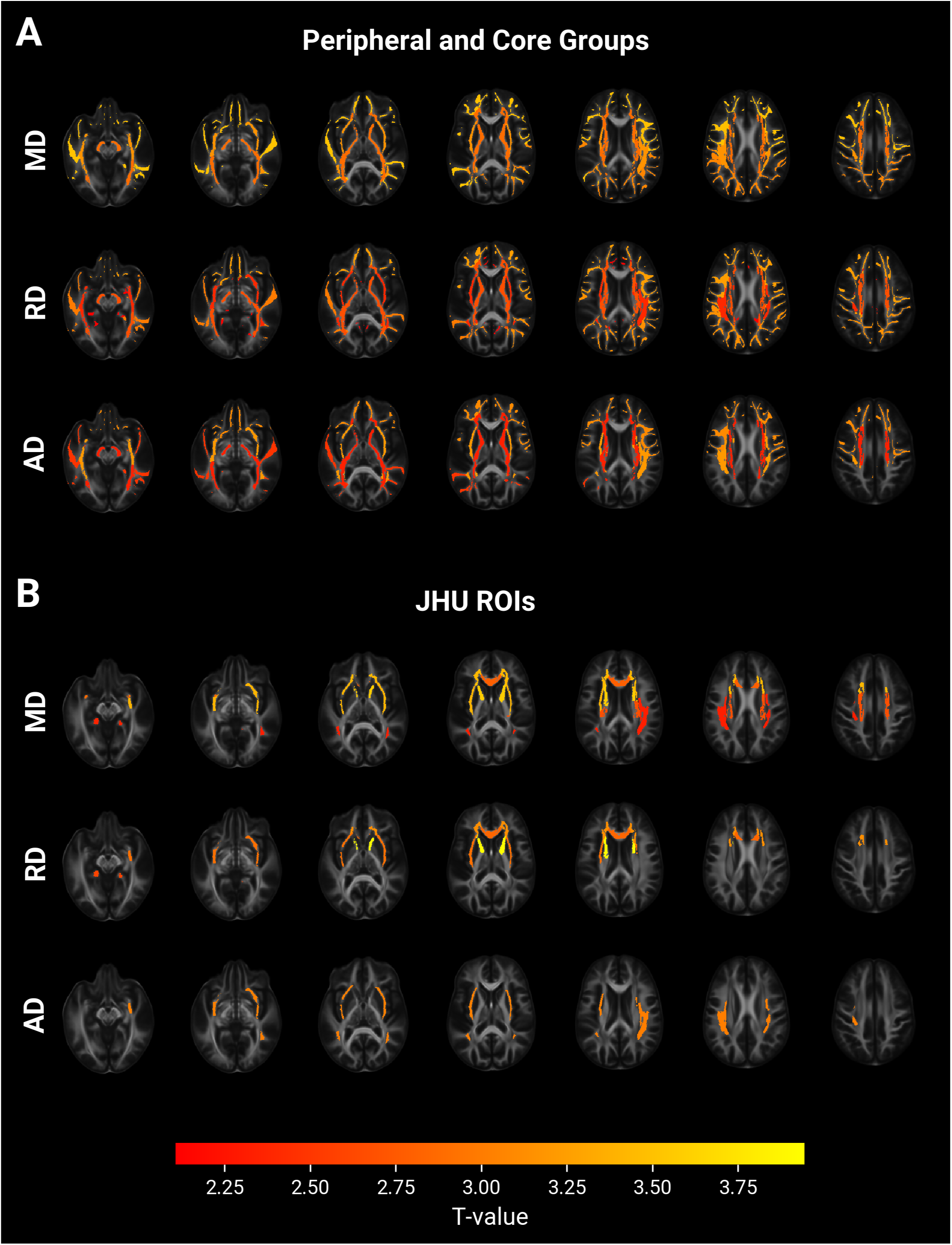
Correlations between microstructural parameters and SANS intercept in the JHSZC dataset. Microstructural measures and SANS intercepts were computed as in Figure 4. Relationships were tested with a linear model with age and sex and covariates. Significant ROIs are coloured according to their T-value. A and B show two nested hierarchical layers, with B at a finer resolution. Comparisons within each layer were corrected for multiple comparisons using FDR. T-values and P-values are shown in Table 5.

**Table 5:** Microstructural measures versus SANS intercept in JHSZC patients.

| Region | $N_{int} \sim -FA$ | | $N_{int} \sim MD$ | | $N_{int} \sim RD$ | | $N_{int} \sim AD$ | |
| --- | --- | --- | --- | --- | --- | --- | --- | --- |
| | $T(24)$ | $P_{corr}$ | $T(24)$ | $P_{corr}$ | $T(24)$ | $P_{corr}$ | $T(24)$ | $P_{corr}$ |
| White Matter | -1.7 | **.048** | 3.2 | **.002** | 2.9 | **.004** | 2.7 | **.006** |
| *Global ROIs* |  |  |  |  |  |  |  |  |
| Core White Matter | -1.9 | .072 | 2.8 | **.005** | 2.6 | **.009** | 2.5 | **.009** |
| Peripheral White Matter | -1.5 | .075 | 3.3 | **.003** | 2.9 | **.007** | 2.7 | **.009** |
| *Regional ROIs* |  |  |  |  |  |  |  |  |
| Association Tracts | -0.9 | .32 | 3.2 | **.008** | 2.4 | **.029** | 3.2 | **.015** |
| Projection Tracts | -2. | .19 | 2.9 | **.01** | 2.7 | **.02** | 2.3 | **.042** |
| Frontal Lobe | -2.4 | .16 | 3.5 | **.006** | 3.2 | **.013** | 3.1 | **.015** |
| Limbic Lobe | -1.4 | .22 | 2. | .053 | 2.1 | **.046** | 1.3 | .13 |
| Posterior Lobe | -1.4 | .22 | 3.1 | **.008** | 3.1 | **.013** | 1.9 | .077 |
| Temporal Lobe | -1.2 | .25 | 3.5 | **.006** | 3. | **.013** | 2.5 | **.04** |
| *Local ROIs* |  |  |  |  |  |  |  |  |
| External capsule | -1.4 | .25 | 3.4 | **.009** | 2.9 | **.027** | 3. | **.036** |
| Inferior fronto-occipital fasciculus | -0.73 | .3 | 2.9 | **.024** | 1.8 | .12 | 2.2 | .1 |
| Superior longitudinal fasciculus | 0.22 | .67 | 2.3 | **.039** | 0.86 | .25 | 3. | **.036** |
| CC Genu | -2.5 | .12 | 2.8 | **.024** | 2.8 | **.027** | 1.9 | .11 |
| Hippocampus | -1.7 | .24 | 2.4 | **.039** | 2.6 | **.038** | 1.3 | .16 |
| Anterior corona radiata | -1.7 | .24 | 3.4 | **.009** | 3.1 | **.027** | 2.5 | .072 |
| Corticospinal tract | -0.33 | .45 | 2.3 | **.039** | 1.5 | .13 | 1.8 | .11 |
| IC Anterior Limb | -3.1 | .056 | 3.6 | **.009** | 3.9 | **.007** | 1.9 | .11 |
| Superior corona radiata | -1.2 | .26 | 2.7 | **.026** | 2.3 | .06 | 1.9 | .11 |

Intercept computed for each subject by fitting a first-order linear model. Statistics computed using unpaired, 1-tailed T-test after regressing age and sex. Bold results indicate significant results (only rows with at least one such result are shown). All p-values corrected using FDR with comparisons in the same hierarchical level.

To further characterize the structural correspondence with symptom scores, the above models were repeated using TBSS to find clusters of significance potentially outside the constraints of the *a priori* ROI parcellation. For MD, RD, and AD, we found extensive clusters with a positive correlation with the SANS intercept, predominately in the bilateral frontal hemisphere peripheral white matter, the genu of the corpus callosum, and the right hemisphere internal and external capsules and corona radiata. We additionally found a negative correlation between FA and the SANS intercept in the genu of the corpus callosom and right hemisphere internal capsule (Figure 7 A). As before, no correlations were found with the SANS slope or with the SAPS intercept or slope.

### 3.5 DTI metrics in TOPSY

We repeated the above analyses in the TOPSY dataset. Many of the patients in the dataset had a large reduction in positive symptoms as measured by the PANSS8-P subscore by their first follow-up visit (Figure 3 B). For the few subjects that had additional follow-ups, their positive symptom scores had a nonlinear trajectory that plateaued in the second session. However, most subjects had only one follow-up, therefore we could not reliably model a nonlinear response in most of our sample. Additional follow-ups were also heavily biased toward patients with minimal negative symptoms (Figure S2). We therefore simplified our analysis by only using the baseline and first follow-up score from each subject.

Because many subjects had the lowest possible score of three in either PANSS8-N or PANSS8-P by their second session, in addition to modelling the score intercept and slope, we additionally split subjects into two groups based on whether they had a subscore of three in their second session, hereon referred to as the “remission” group. This was done separately for each subscore. We then tested for differences in DTI metrics between these groups. Models and 1-tailed contrasts were used as with the JHSZC dataset. For the additional group split, we tested if FA was higher in the recovery group, and if MD, RD, and AD were lower.

Significantly higher global FA was found in the PANSS8-N remission group (Figure 6 A). The finding was replicated in both the core and peripheral white matter following FDR correction (Figure 6 B). Finally, we observed this result in the frontal and posterior lobe ROIs, and in the limbic tract grouping of JHU atlas ROIs (Figure 6 C). No effect on FA was found for the PANSS8-N slope or intercept, or for any of the PANSS8-P metrics. No effects were found for any other DTI parameter.

**Figure 6:**
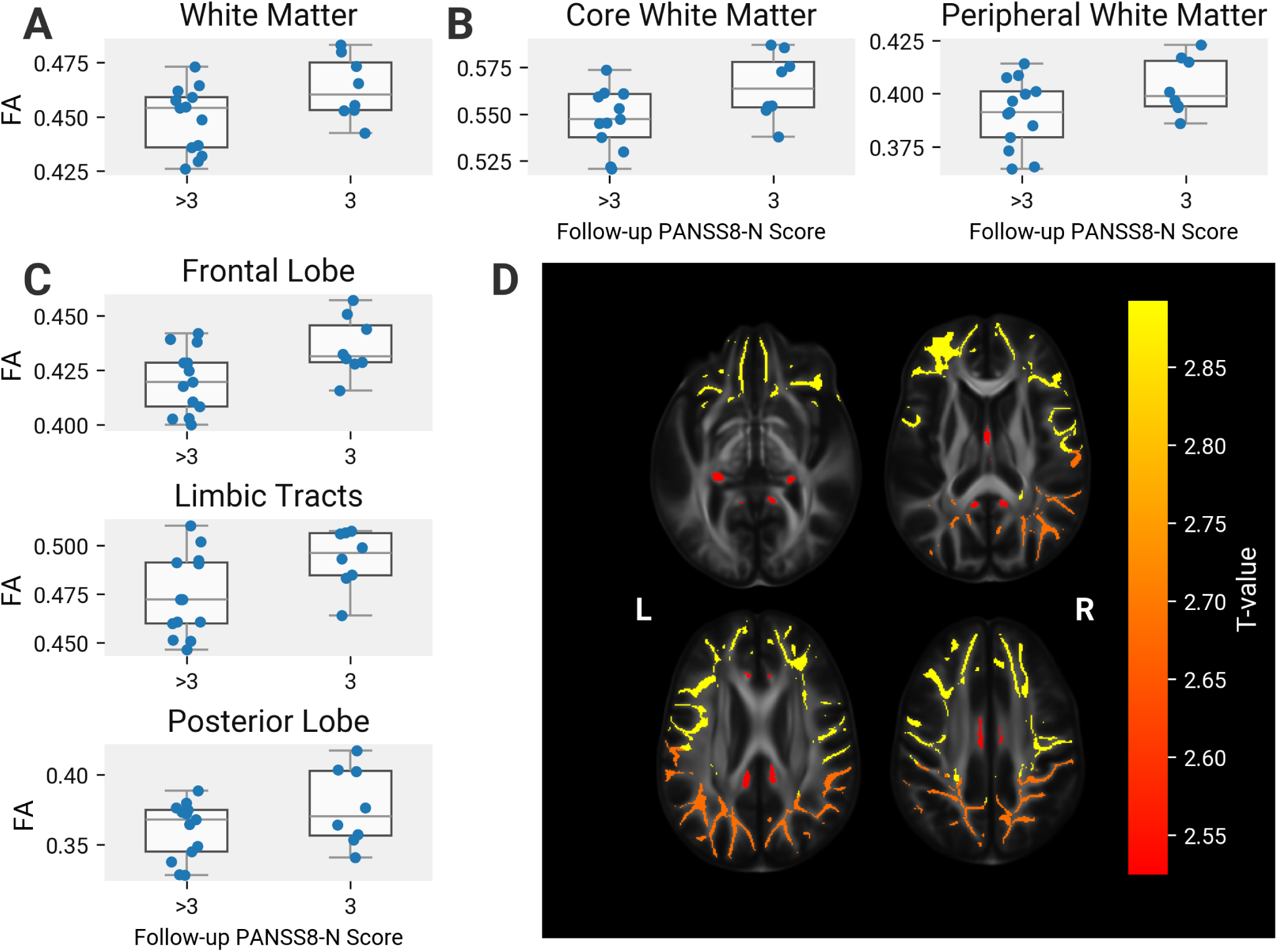
Correlations between microstructural parameters and the PANSS8-N follow-up score. Microstructural measures are computed as in Figure 4. Subjects are grouped based on whether their PANSS8-N score at their follow-up session was equal to 3, the lowest possible score. Relationships were tested with a linear model with age and sex and covariates. All comparisons shown are signficant. A, B, C: ROIs from different nested hierarchical layers at successively higher resolutions. D: the location of the regions in C with their T-values. Comparisons within each layer were corrected for multiple comparisons using FDR. T-values and P-values are shown in Table 6.

**Table 6:** DTI measures in TOPSY patients with a follow-up PANSS8-N score of 3 versus higher than 3.

| Region | $N_{int} \sim -FA$ | | $N_{int} \sim MD$ | | $N_{int} \sim RD$ | | $N_{int} \sim AD$ | |
| --- | --- | --- | --- | --- | --- | --- | --- | --- |
| | $T(17)$ | $P_{corr}$ | $T(17)$ | $P_{corr}$ | $T(17)$ | $P_{corr}$ | $T(17)$ | $P_{corr}$ |
| White Matter | 2.6 | **.01** | -1.1 | .14 | -1.6 | .059 | 0.41 | .66 |
| *Global ROIs* |  |  |  |  |  |  |  |  |
| Core White Matter | 2.2 | **.021** | -0.62 | .27 | -1.5 | .077 | 2. | .97 |
| Peripheral White Matter | 2.6 | **.019** | -1.4 | .2 | -1.7 | .077 | -0.4 | .69 |
| *Regional ROIs* |  |  |  |  |  |  |  |  |
| Limbic Tracts | 2.5 | **.044** | -0.92 | .37 | -1.9 | .18 | 1.8 | 1.0 |
| Frontal Lobe | 2.9 | **.044** | -1.3 | .35 | -1.8 | .18 | -0.28 | .94 |
| Posterior Lobe | 2.7 | **.044** | -1.1 | .35 | -1.5 | .21 | -0.31 | .94 |

Statistics computed using paired, 1-tailed T-tests after regressing age and sex. Bold results indicate significant results (only rows with at least one such result are shown). All p-values corrected using FDR with comparisons in the same hierarchical level.

**Table 7:** Subnetworks with a significant association between DTI parameter and a follow-up PANSS8-N score of 3.

| Param | # Connections | $P_{corr}$ |
| --- | --- | --- |
| RD | 486 | .029 |
| FA | 1278 | .007 |

As with JHSZC, we used TBSS to find data-driven clusters of significantly affected regions. Clusters of significantly higher FA in the PANSS8-N remission group were found bilaterally in the parietal peripheral white matter, occipital peripheral white matter, splenium of the corpus callosum, left hemisphere internal capsule, and bilateral superior longitudinal fasciculus (Figure 7 B). No other significant clusters were found.

**Figure 7:**
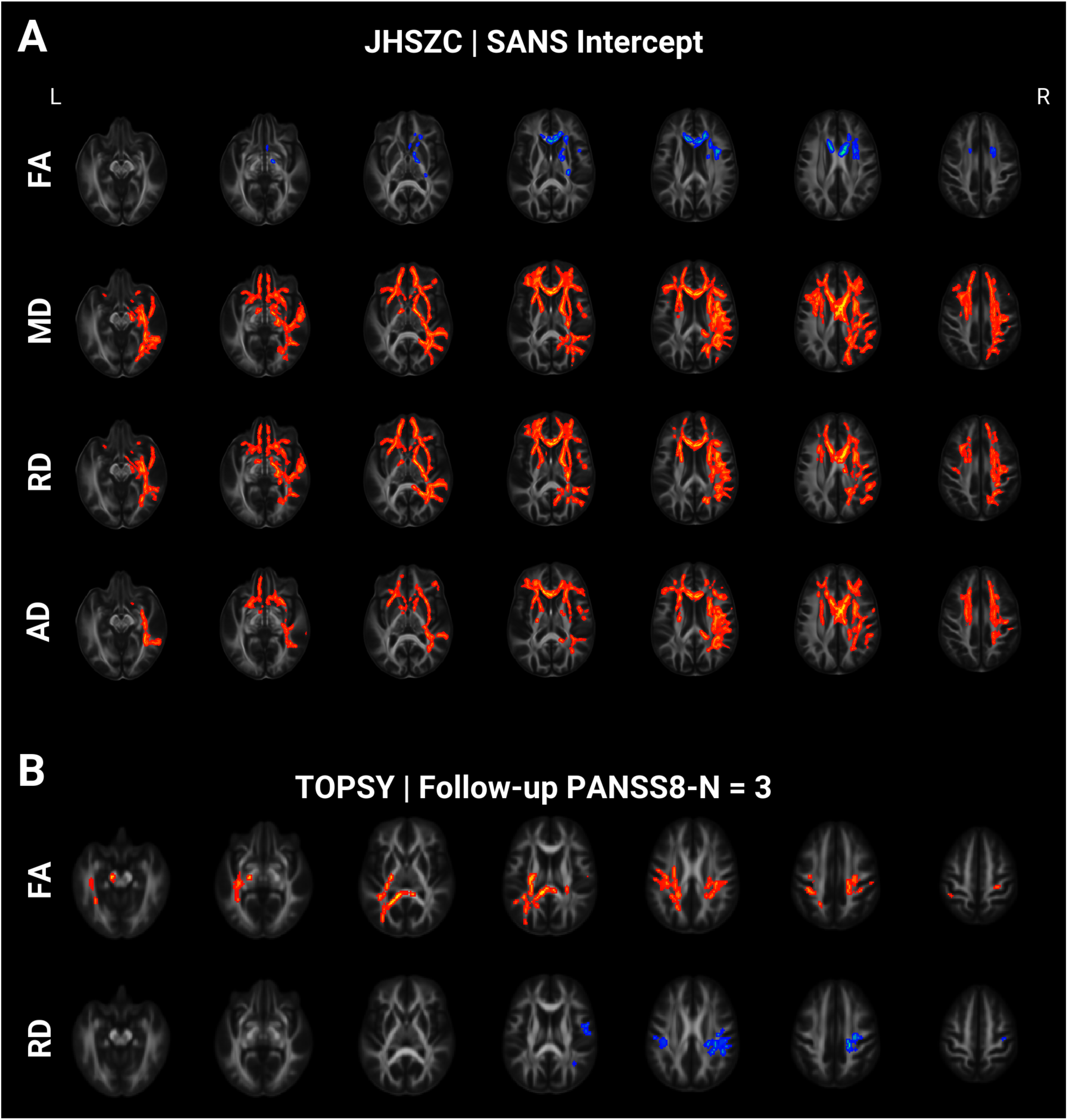
Regions associated with negative symptoms. Displayed clusters significantly correlate with the metric of interest as determined using TFCE (10,000 samples, FWER < 0.05). Clusters are localized to the TBSS-derived FA skeleton and inflated for visualization. A: Measures in JHSCZ patients compared with the SANS intercept, as described in Figure 4. B: Effect of PANSS8-N recovery, as described in Figure 6, on microstructure in TOPSY patients.

Because the TOPSY diffusion data was acquired using a high angular-resolution diffusion imaging (HARDI) diffusion paradigm, we were able to perform robust probabilistic tractography. We sampled the DTI parameters along the generated streamlines and generated weighted structural connectomes using the Brainnetome atlas. We then fit the above models on the weighted connectomes, performing statistics with NBS. Confirming our TBSS results, we found a large subnetwork connecting ROIs across both hemispheres with significantly higher FA in the PANSS8-N remission group. We also found a subnetwork with significantly lower RD in the same group, predominately within the right hemisphere parietal lobe and between the right hemisphere frontal and parietal lobes (Figure 8). No other significant subnetworks were found.

**Figure 8:**
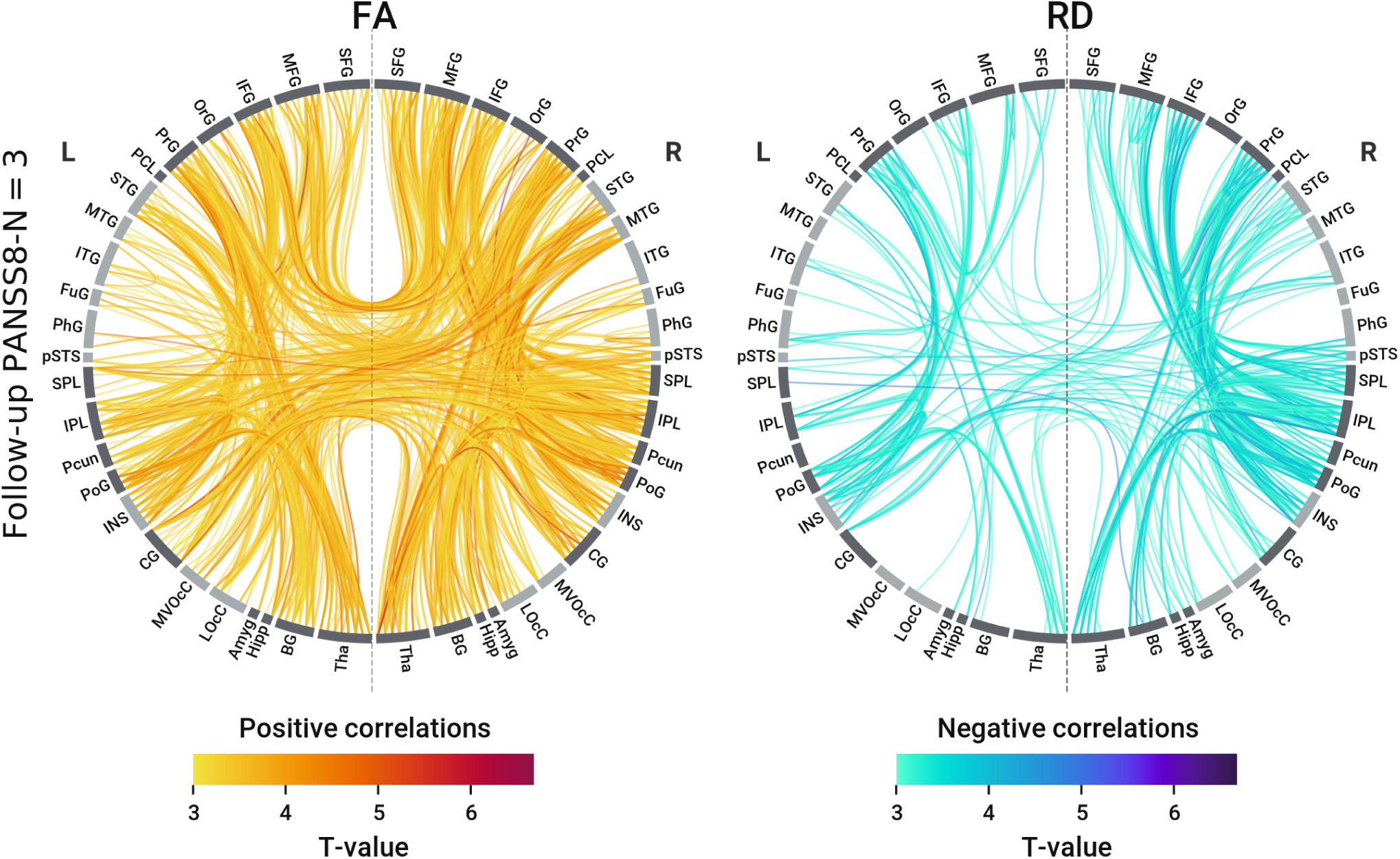
Networks associated with negative symptoms in the TOPSY dataset. Microstructural measures are averaged across sessions. Values for each connection were measured by sampling along the constituent streamlines. The recovery score is computed as in Figure 6. In each network diagram, lines represent connections significantly correlated with the corresponding PANSS8-N derivative measure as determined using NBS (10,000 samples, 𝑇_𝑡ℎ𝑟𝑒𝑠ℎ_ = 3, FWER < 0.05). Left diagram represents connections with significantly higher FA in patients a PANSS8-N score of 3 (the lowest possible score) a their follow-up session. Right hemisphere is the same, but with lower RD as the correlate. Gyral abbreviations are given in Table 1. Subnetwork size and p-values are given in Table 7.

## 4 Discussion

In this study, we failed to find significant longitudinal effects in DTI parameters in patients from two different datasets obtained at different field strengths in the same phase of illness (3 years since diagnosis). Thus, our results do not provide evidence for the neurodegenerative hypothesis for white matter in psychosis. As noted previously, FA in patients putatively reaches an early developmental peak in their twenties. If any longitudinal differences (i.e. deviations from the expected slope) are to be found between patients and controls, we would reasonably expect them to appear within the timeframe studied.

Nonetheless, some alternative explanations may yet explain our results. First, our patients may have been too young for neurodegenerative white matter changes to occur, as disease-driven decline may not manifest until closer to the developmental peak at around thirty years of age (15).

Second, our study may not have been long enough to detect FA decline, which normally ranges from 0.1 to 0.5% annualized change (68). We note, however, that increased follow-up time will not necessarily increase experimental power. FA in patients cannot perpetually decline faster than HCs, as this would result in implausibly low FA in old age. Thus, rates of change in the two groups must eventually equilibrate as FA reaches its lower plateau. It is not clear when this equilibration might happen, as quadratic models fit to cross-sectional data tend to give unreliable rate estimates (69).

Third, our sample size may have been too small. A larger sample may have detected more subtle effects, however, the clinical relevance of such small differences seen in larger datasets may be questionable.

Although no prominent group differences were observed, we did find associations between DTI and negative symptoms in both datasets, consistent with our prior observations (1). In the JHSZC dataset, all four DTI parameters had extensive interactions with the SANS intercept. The absence of a correlation with the SANS slope may relate to the relative lack of variation in subject scores across the JHSZC dataset sessions, especially as these patients were more established on their treatments than the TOPSY cohort. The fact that the intercept, rather than slope, correlated with DTI parameters suggests these associations are not related to medication and treatment response or recovery.

Negative symptoms are often secondary to, and ameliorate upon treatment of, positive symptoms (70). In our TOPSY dataset, where positive symptoms in most patients responded well to treatment and depressive episodes were ruled out via clinical consensus, residual negative symptoms in the second session may represent persisting, primary symptoms. If so, our treatment response grouping distinguished patients with and without primary negative symptoms, finding more chronic-like DTI parameters in the primary symptom group. Previous studies have in fact shown that patients with deficit schizophrenia, defined by the prominence of primary negative symptoms, have lower FA than non-deficit patients (71–75) (but see in contrast (76)).

### 4.1 Limitations

In the JHSZC dataset, patients were not age or sex matched to controls. Age, in particular, is known to affect the rate of change for DTI parameters. Analysis with an age and sex-matched subset of the dataset also failed to find different longitudinal trajectories, but this may have been affected by the smaller sample size. Longitudinal designs are subject to attrition bias; for instance, more severely ill patients may be less likely to remain in the study (although we found no association between drop-out and clinical scores). Additionally, our investigations of clinical symptoms were exploratory, and thus used liberal significance thresholds. FDR was used for multiple comparisons in our ROI analysis rather than a family-wise error rate method, and no correcting was done across different parameter-clinical score combinations.

### 4.2 Conclusion

In summary, although we do not find evidence for longitudinal change of white matter integrity in the first few years of psychosis, we did observe complementary associations between white matter state and negative symptoms in two completely independent datasets, each of which drew patients from different geographic regions, and used different acquisition protocols and clinical scales. Thus, our data fail to support a neurodegenerative hypothesis, but do support a mediating effect of white matter on illness severity, especially on negative symptom burden.

## Supporting information

JHSZC Longitudinal Stats

JHSZC Age-matched Longitudinal Stats

JHSZC Symptom Score Stats

TOPSY Longitudinal Stats

TOPSY Symptom Score Stats

## Acknowledgements

Special thanks to Ravi Menon for his leadership for the imaging component of TOPSY cohort. We also thank Peter Williamson, Jean Theberge, Robert Bartha, Robert Hegele, Justin Hicks, Richard Neufeld and Keith St. Lawrence for their assistance in funding acquisition for this study.

## References

1. Van Dyken PC, MacKinley M, Khan AR, Palaniyappan L (2024): Cortical Network Disruption Is Minimal in Early Stages of Psychosis. Schizophrenia Bulletin Open 5: sgae010.

2. Deng Y, Hung KSY, Lui SSY, Chui WWH, Lee JCW, Wang Y, et al. (2019): Tractography-based classification in distinguishing patients with first-episode schizophrenia from healthy individuals. Progress in Neuro-Psychopharmacology & Biological Psychiatry 88: 66–73.

3. Filippi M, Canu E, Gasparotti R, Agosta F, Valsecchi P, Lodoli G, et al. (2014): Patterns of Brain Structural Changes in First-Contact, Antipsychotic Drug-Naïve Patients with Schizophrenia. American Journal of Neuroradiology 35: 30–37.

4. Kraguljac NV, Anthony T, Morgan CJ, Jindal RD, Burger MS, Lahti AC (2021): White Matter Integrity, Duration of Untreated Psychosis, and Antipsychotic Treatment Response in Medication-Naïve First-Episode Psychosis Patients. Molecular Psychiatry 26: 5347–5356.

5. Keymer-Gausset A, Alonso-Solís A, Corripio I, Sauras-Quetcuti RB, Pomarol-Clotet E, Canales-Rodriguez EJ, et al. (2018): Gray and white matter changes and their relation to illness trajectory in first episode psychosis. European Neuropsychopharmacology: The Journal of the European College of Neuropsychopharmacology 28: 392–400.

6. Melicher T, Horacek J, Hlinka J, Spaniel F, Tintera J, Ibrahim I, et al. (2015): White matter changes in first episode psychosis and their relation to the size of sample studied: a DTI study. Schizophrenia Research 162: 22–28.

7. Maximo JO, Kraguljac NV, Rountree BG, Lahti AC (2021): Structural and Functional Default Mode Network Connectivity and Antipsychotic Treatment Response in Medication-Naïve First Episode Psychosis Patients. Schizophrenia Bulletin Open 2: sgab032.

8. Pérez-Iglesias R, Tordesillas-Gutiérrez D, Barker GJ, McGuire PK, Roiz-Santiañez R, Mata I, et al. (2010): White matter defects in first episode psychosis patients: a voxelwise analysis of diffusion tensor imaging. NeuroImage 49: 199–204.

9. Ruef A, Curtis L, Moy G, Bessero S, Badan Bâ M, Lazeyras F, et al. (2012): Magnetic resonance imaging correlates of first-episode psychosis in young adult male patients: combined analysis of grey and white matter. Journal of psychiatry & neuroscience: JPN 37: 305–312.

10. Serpa MH, Doshi J, Erus G, Chaim-Avancini TM, Cavallet M, Bilt MT van de, et al. (2017): State-dependent microstructural white matter changes in drug-naïve patients with first-episode psychosis. Psychological Medicine 47: 2613–2627.

11. Faria AV, Zhao Y, Ye C, Hsu J, Yang K, Cifuentes E, et al. (2021): Multimodal MRI Assessment for First Episode Psychosis: A Major Change in the Thalamus and an Efficient Stratification of a Subgroup. Human Brain Mapping 42: 1034–1053.

12. Szeszko PR, Ardekani BA, Ashtari M, Kumra S, Robinson DG, Sevy S, et al. (2005): White matter abnormalities in first-episode schizophrenia or schizoaffective disorder: a diffusion tensor imaging study. The American Journal of Psychiatry 162: 602–605.

13. Lee DY, Smith GN, Su W, Honer WG, Macewan GW, Lapointe JS, et al. (2012): White matter tract abnormalities in first-episode psychosis. Schizophrenia Research 141: 29–34.

14. Alvarado-Alanis P, León-Ortiz P, Reyes-Madrigal F, Favila R, Rodríguez-Mayoral O, Nicolini H, et al. (2015): Abnormal white matter integrity in antipsychotic-naïve first-episode psychosis patients assessed by a DTI principal component analysis. Schizophrenia Research 162: 14–21.

15. Cetin-Karayumak S, Di Biase MA, Chunga N, Reid B, Somes N, Lyall AE, et al. (2020): White Matter Abnormalities Across the Lifespan of Schizophrenia: A Harmonized Multi-Site Diffusion MRI Study. Molecular Psychiatry 25: 3208–3219.

16. Kochunov P, Ganjgahi H, Winkler A, Kelly S, Shukla DK, Du X, et al. (2016): Heterochronicity of White Matter Development and Aging Explains Regional Patient Control Differences in Schizophrenia. Human Brain Mapping 37: 4673–4688.

17. Kochunov P, Glahn DC, Rowland LM, Olvera RL, Winkler A, Yang Y-H, et al. (2013): Testing the Hypothesis of Accelerated Cerebral White Matter Aging in Schizophrenia and Major Depression. Biological Psychiatry 73: 482–491.

18. Wright SN, Kochunov P, Chiappelli J, McMahon RP, Muellerklein F, Wijtenburg SA, et al. (2014): Accelerated white matter aging in schizophrenia: Role of white matter blood perfusion. Neurobiology of Aging 35: 2411–2418.

19. Tronchin G, McPhilemy G, Ahmed M, Kilmartin L, Costello L, Forde NJ, et al. (2021): White matter microstructure and structural networks in treatment-resistant schizophrenia patients after commencing clozapine treatment: A longitudinal diffusion imaging study. Psychiatry Research 298: 113772.

20. Domen P, Peeters S, Michielse S, Gronenschild E, Viechtbauer W, Roebroeck A, et al. (2017): Differential Time Course of Microstructural White Matter in Patients With Psychotic Disorder and Individuals at Risk: A 3-Year Follow-up Study. Schizophrenia Bulletin 43: 160–170.

21. Bergé D, Mané A, Lesh TA, Bioque M, Barcones F, Gonzalez-Pinto AM, et al. (2020): Elevated Extracellular Free-Water in a Multicentric First-Episode Psychosis Sample, Decrease During the First 2 Years of Illness. Schizophrenia Bulletin 46: 846–856.

22. Mittal VA, Dean DJ, Bernard JA, Orr JM, Pelletier-Baldelli A, Carol EE, et al. (2014): Neurological Soft Signs Predict Abnormal Cerebellar-Thalamic Tract Development and Negative Symptoms in Adolescents at High Risk for Psychosis: A Longitudinal Perspective. Schizophrenia Bulletin 40: 1204–1215.

23. Bernard JA, Orr JM, Mittal VA (2015): Abnormal Hippocampal–Thalamic White Matter Tract Development and Positive Symptom Course in Individuals at Ultra-High Risk for Psychosis. npj Schizophrenia 1: 1–6.

24. Kristensen TD, Glenthøj LB, Raghava JM, Syeda W, Mandl RCW, Wenneberg C, et al. (2021): Changes in negative symptoms are linked to white matter changes in superior longitudinal fasciculus in individuals at ultra-high risk for psychosis. Schizophrenia Research 237: 192–201.

25. Roalf DR, Garza AG de la, Rosen A, Calkins ME, Moore TM, Quarmley M, et al. (2020): Alterations in White Matter Microstructure in Individuals at Persistent Risk for Psychosis. Molecular Psychiatry 25: 2441–2454.

26. Krakauer K, Nordentoft M, Glenthøj BY, Raghava JM, Nordholm D, Randers L, et al. (2018): White matter maturation during 12 months in individuals at ultra-high-risk for psychosis. Acta Psychiatrica Scandinavica 137: 65–78.

27. Bernard JA, Orr JM, Mittal VA (2017): Cerebello-thalamo-cortical networks predict positive symptom progression in individuals at ultra-high risk for psychosis. NeuroImage: Clinical 14: 622–628.

28. Beck D, Lange A-MG de, Maximov II, Richard G, Andreassen OA, Nordvik JE, Westlye LT (2021): White Matter Microstructure Across the Adult Lifespan: A Mixed Longitudinal and Cross-Sectional Study Using Advanced Diffusion Models and Brain-Age Prediction. NeuroImage 224: 117441.

29. Kochunov P, Williamson DE, Lancaster J, Fox P, Cornell J, Blangero J, Glahn DC (2012): Fractional anisotropy of water diffusion in cerebral white matter across the lifespan. Neurobiology of Aging 33: 9–20.

30. Schilling KG, Archer D, Rheault F, Lyu I, Huo Y, Cai LY, et al. (2023): Superficial White Mat- ter Across Development, Young Adulthood, and Aging: Volume, Thickness, and Relationship with Cortical Features. Brain Structure and Function 228: 1019–1031.

31. Sexton CE, Walhovd KB, Storsve AB, Tamnes CK, Westlye LT, Johansen-Berg H, Fjell AM (2014): Accelerated Changes in White Matter Microstructure During Aging: A Longitudinal Diffusion Tensor Imaging Study. Journal of Neuroscience 34: 15425–15436.

32. Kochunov P, Hong LE (2014): Neurodevelopmental and Neurodegenerative Models of Schizophrenia: White Matter at the Center Stage. Schizophrenia Bulletin 40: 721–728.

33. Kelly S, Jahanshad N, Zalesky A, Kochunov P, Agartz I, Alloza C, et al. (2018): Widespread White Matter Microstructural Differences in Schizophrenia Across 4322 Individuals: Results from the ENIGMA Schizophrenia DTI Working Group. Molecular Psychiatry 23: 1261–1269.

34. Chen F, Mihaljevic M, Hou Z, Li Y, Lu H, Mori S, et al. (2023): Relation between white matter integrity, perfusion, and processing speed in early-stage schizophrenia. Journal of Psychiatric Research 163: 166–171.

35. Faria AV, Zhao Y, Ye C, Hsu J, Yang K, Cifuentes E, et al. (2020): Multimodal MRI assessment for first episode psychosis: A major change in the thalamus and an efficient stratification of a subgroup. Human Brain Mapping 42: 1034–1053.

36. Faria AV, Crawford J, Ye C, Hsu J, Kenkare A, Schretlen D, Sawa A (2019): Relationship between neuropsychological behavior and brain white matter in first-episode psychosis. Schizophrenia Research 208: 49–54.

37. Leckman JF, Sholomskas D, Thompson WD, Belanger A, Weissman MM (1982): Best estimate of lifetime psychiatric diagnosis: a methodological study. Archives of General Psychiatry 39: 879–883.

38. Marques JP, Kober T, Krueger G, Zwaag W van der, Van de Moortele P-F, Gruetter R (2010): MP2RAGE, a self bias-field corrected sequence for improved segmentation and T1-mapping at high field. NeuroImage 49: 1271–1281.

39. Fan L, Li H, Zhuo J, Zhang Y, Wang J, Chen L, et al. (2016): The Human Brainnetome Atlas: A New Brain Atlas Based on Connectional Architecture. Cerebral Cortex 26: 3508–3526.

40. Iglesias JE, Billot B, Balbastre Y, Tabari A, Conklin J, Gilberto González R, et al. (2021): Joint super-resolution and synthesis of 1 mm isotropic MP-RAGE volumes from clinical MRI exams with scans of different orientation, resolution and contrast. NeuroImage 237: 118206.

41. Avants BB, Epstein CL, Grossman M, Gee JC (2008): Symmetric diffeomorphic image registration with cross-correlation: evaluating automated labeling of elderly and neurodegenerative brain. Medical Image Analysis 12: 26–41.

42. Benjamini Y, Hochberg Y (1995): Controlling the false discovery rate: A practical and powerful approach to multiple testing. Journal of the Royal statistical society: series B (Methodological) 57: 289–300.

43. Simmonds DJ, Hallquist MN, Asato M, Luna B (2014): Developmental stages and sex differences of white matter and behavioral development through adolescence: A longitudinal diffusion tensor imaging (DTI) study. NeuroImage 92: 356–368.

44. Team RC (2024): R: A Language and Environment for Statistical Computing. Vienna, Austria: R Foundation for Statistical Computing. Retrieved from https://www.R-project.org/

45. Bates D, Mächler M, Bolker B, Walker S (2015): Fitting linear mixed-effects models using Lme4. Journal of Statistical Software 67: 1–48.

46. Kuznetsova A, Brockhoff PB, Christensen RHB (2017): lmerTest package: Tests in linear mixed effects models. Journal of Statistical Software 82: 1–26.

47. Satterthwaite FE (1946): An Approximate Distribution of Estimates of Variance Components. Biometrics Bulletin 2: 110–114.

48. Winkler AM, Ridgway GR, Webster MA, Smith SM, Nichols TE (2014): Permutation Inference for the General Linear Model. NeuroImage 92: 381–397.

49. Zalesky A, Fornito A, Bullmore ET (2010): Network-based statistic: identifying differences in brain networks. NeuroImage 53: 1197–1207.

50. Harris CR, Millman KJ, Walt SJ van der, Gommers R, Virtanen P, Cournapeau D, et al. (2020): Array Programming with NumPy. Nature 585: 357–362.

51. McKinney W (2010): Data Structures for Statistical Computing in Python. 56–61.

52. Reback J, jbrockmendel, McKinney W, Bossche JV den, Augspurger T, Cloud P, et al. (2022, February 12): Pandas-dev/pandas: Pandas 1.4.1. Zenodo. 10.5281/zenodo.60532 72

53. Hoyer S, Hamman J (2017): Xarray: N-D labeled Arrays and Datasets in Python. 5: 10.

54. Hoyer S, Roos M, Joseph H, Magin J, Cherian D, Fitzgerald C, et al. (2022, July 22): Xarray. Zenodo. 10.5281/zenodo.6885151

55. Virtanen P, Gommers R, Oliphant TE, Haberland M, Reddy T, Cournapeau D, et al. (2020): SciPy 1.0: Fundamental algorithms for scientific computing in python. Nature Methods 17: 261–272.

56. Seabold S, Perktold J (2010): Statsmodels: Econometric and statistical modeling with python. 9th Python in Science Conference. 10.25080/Majora-92bf1922-011

57. Hagberg AA, Schult DA, Swart PJ (2008): Exploring network structure, dynamics, and function using NetworkX. In: Varoquaux G, Vaught T, Millman J, editors. Proceedings of the 7th Python in Science Conference. Pasadena, CA USA, pp 11–15.

58. Waskom ML (2021): Seaborn: Statistical data visualization. Journal of Open Source Software 6: 3021.

59. Hunter JD (2007): Matplotlib: A 2D graphics environment. Computing in Science & Engineering 9: 90–95.

60. Ciric R, Thompson WH, Lorenz R, Goncalves M, MacNicol EE, Markiewicz CJ, et al. (2022): TemplateFlow: FAIR-sharing of Multi-Scale, Multi-Species Brain Models. Nature Methods 19: 1568– 1571.

61. Peixoto TP (2014): The graph-tool python library. *figshare*. 10.6084/m9.figshare.1164194

62. Yarkoni T, Markiewicz C, Vega A de la, Gorgolewski K, Salo T, Halchenko Y, et al. (2019): PyBIDS: Python tools for BIDS datasets. Journal of Open Source Software 4: 1294.

63. Yarkoni T, Markiewicz CJ, Vega A de la, Gorgolewski KJ, Salo T, Halchenko YO, et al. (2023, August 16): PyBIDS: Python tools for BIDS datasets. Zenodo. 10.5281/zenodo.8253830

64. Brett M, Markiewicz CJ, Hanke M, Côté M-A, Cipollini B, McCarthy P, et al. (2020, November 28): Nipy/nibabel: 3.2.1. Zenodo. 10.5281/zenodo.4295521

65. Contributors N, Chamma A, Frau-Pascual A, Rothberg A, Abadie A, Abraham A, et al. (2023, October 1): Nilearn. Zenodo. 10.5281/zenodo.8397157

66. Preedy VR (2016): Handbook of Cannabis and Related Pathologies: Biology, Pharmacology, Diagnosis, and Treatment. Academic Press.

67. Goldman HH, Skodol AE, Lave TR (1992): Revising axis V for DSM-IV: a review of measures of social functioning. The American Journal of Psychiatry 149: 1148–1156.

68. Schilling KG, Archer D, Yeh F-C, Rheault F, Cai LY, Shafer A, et al. (2023): Short superficial white matter and aging: A longitudinal multi-site study of 1293 subjects and 2711 sessions. Aging Brain 3: 100067.

69. Fjell AndersM, Walhovd KB, Westlye LT, Østby Y, Tamnes CK, Jernigan TL, et al. (2010): When does brain aging accelerate? Dangers of quadratic fits in cross-sectional studies. NeuroImage 50: 1376–1383.

70. Lutgens D, Joober R, Iyer S, Lepage M, Norman R, Schmitz N, et al. (2019): Progress of negative symptoms over the initial 5 years of a first episode of psychosis. Psychological Medicine 49: 66–74.

71. Lei W, Li N, Deng W, Li M, Huang C, Ma X, et al. (2015): White Matter Alterations in First Episode Treatment-Naïve Patients with Deficit Schizophrenia: A Combined VBM and DTI Study. Scientific Reports 5: 12994.

72. Rowland LM, Spieker EA, Francis A, Barker PB, Carpenter WT, Buchanan RW (2009): White Matter Alterations in Deficit Schizophrenia. Neuropsychopharmacology 34: 1514–1522.

73. Voineskos AN, Foussias G, Lerch J, Felsky D, Remington G, Rajji TK, et al. (2013): Neuroimaging Evidence for the Deficit Subtype of Schizophrenia. JAMA Psychiatry 70: 472–480.

74. Sun H, Lui S, Yao L, Deng W, Xiao Y, Zhang W, et al. (2015): Two Patterns of White Matter Abnormalities in Medication-Naive Patients With First-Episode Schizophrenia Revealed by Diffusion Tensor Imaging and Cluster Analysis. JAMA Psychiatry 72: 678–686.

75. Smirnova LP, Yarnykh VL, Parshukova DA, Kornetova EG, Semke AV, Usova AV, et al. (2021): Global Hypomyelination of the Brain White and Gray Matter in Schizophrenia: Quantitative Imaging Using Macromolecular Proton Fraction. Translational Psychiatry 11: 1–7.

76. Spalletta G, De Rossi P, Piras F, Iorio M, Dacquino C, Scanu F, et al. (2015): Brain white matter microstructure in deficit and non-deficit subtypes of schizophrenia. Psychiatry Research: Neuroimaging 231: 252–261.

