## Supplementary Data for "Stable White Matter Structure in the First Three Years after Psychosis Onset"

### 1 Supplementary Figures

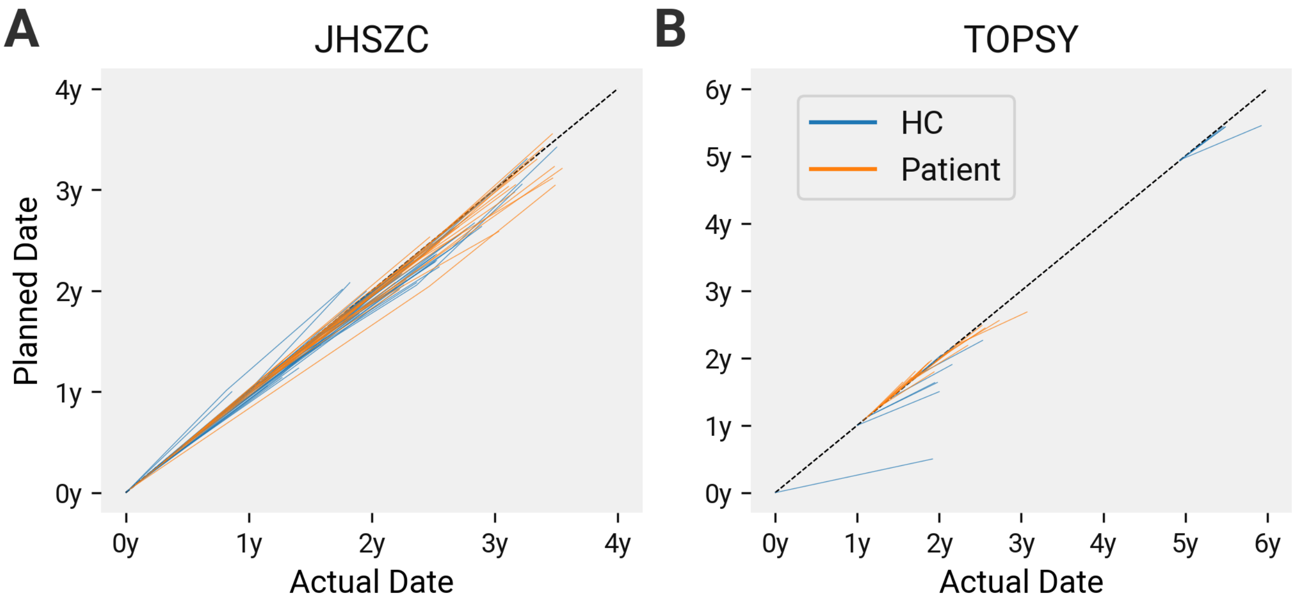


Figure S1: Actual scan dates versus target dates. Individual line segments connect the scan dates from individual subjects. Dates are given as time relative to study onset. Dashed, black line shows the expected slope of subjects scanned at the protocol-specified frequency: 1yr for JHSZC and 6 months for TOPSY. Shallower slopes reflect longer than expected scan-scan intervals; steeper represent shorter intervals.

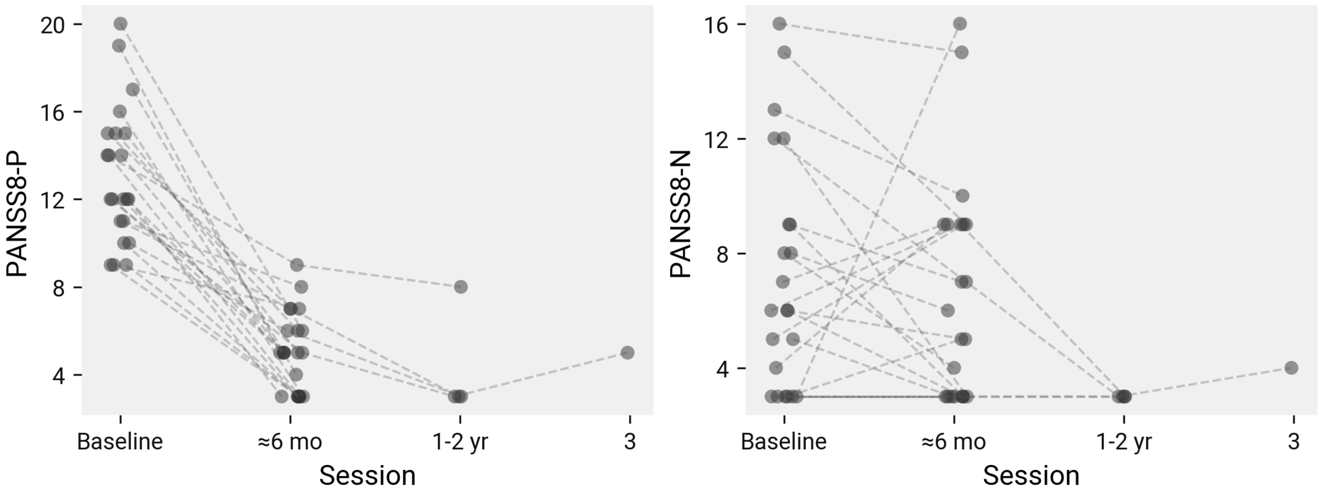


Figure S2: Clinical scores from TOPSY patients across all sessions. Each dashed line corresponds to a different subject.

### 2 Supplementary Tables

Table S1: JHSZC demographic statistics: HC vs patient.

| | Baseline | 1yr | 2yr |
|---|---|---|---|
| Sex (M/F) | $\chi^2(1) = 3.59$, $P = .058$ | $\chi^2(1) = 3.33$, $P = .068$ | $\chi^2(1) = 4.4$, $\boldsymbol{P = .036}$ |
| Age | $T(64) = 2.22$, $\boldsymbol{P = .03}$ | $T(60) = 2.48$, $\boldsymbol{P = .016}$ | $T(29) = 3.01$, $\boldsymbol{P = .0053}$ |
| Ethnicity (B/EA/O/W) | $\chi^2(3) = 2.36$, $P = .5$ | $\chi^2(3) = 3.43$, $P = .33$ | $\chi^2(3) = 2.25$, $P = .52$ |
| Handedness (R/L) | $\chi^2(1) = 0.00947$, $P = .92$ | $\chi^2(1) = 0.14$, $P = .71$ | $\chi^2(1) = 0.0887$, $P = .77$ |
| Smoker (Yes/No) | $\chi^2(1) = 1.56$, $P = .21$ | $\chi^2(1) = 8.92$, $\boldsymbol{P = .0028}$ | $\chi^2(1) = 3.34$, $P = .068$ |
| Cannabis (Yes/No) | $\chi^2(1) = 0.629$, $P = .43$ | $\chi^2(1) = 0.972$, $P = .32$ | $\chi^2(1) = 0.252$, $P = .62$ |

B=Black/African; EA=East Asian; O=Other/Unknown; W=White/European

Table S2: TOPSY demographic statistics.

| | Baseline | 6 mo |
|---|---|---|
| Sex (M/F) | $\chi^2(1) = 0.625$, $P = .43$ | $\chi^2(1) = 0.625$, $P = .43$ |
| Age | $T(32) = -0.508$, $P = .61$ | $T(32) = -0.323$, $P = .75$ |
| Handedness (R/L/A) | $\chi^2(1) = 0$, $P = 1$ | $\chi^2(1) = 0$, $P = 1$ |
| Education | $T(32) = 2.39$, $\boldsymbol{P = .023}$ | $T(32) = 2.44$, $\boldsymbol{P = .02}$ |
| SES | $T(31) = -1.49$, $P = .15$ | $T(31) = -1.49$, $P = .15$ |
| CAST | $T(29) = -2.53$, $\boldsymbol{P = .017}$ | |
| AUDIT-C | $T(27) = -0.794$, $P = .43$ | |
| Smoker (yes/no) | $\chi^2(1) = 2.73$, $P = .098$ | $\chi^2(1) = 0.0145$, $P = .9$ |
| Cannabis (yes/no) | $\chi^2(1) = 1.91$, $P = .17$ | $\chi^2(1) = 0$, $P = 1$ |
| SOFAS | $T(30) = 9.8$, $\boldsymbol{P < .001}$ | $T(24) = 3.11$, $\boldsymbol{P = .0048}$ |

B=Black/African; C=Caribbean/North American Black; EA=East Asian; W=White/European; CAST=Cannabis Abuse Screening Test; SES=Socioeconomic status; AUDIT-C=Alcohol Use Disorders Identification Test

Table S3: JHSZC dropout demographics.

| | Healthy Control | | | Early Psychosis | | |
|---|---|---|---|---|---|---|
| | Dropout (n=54) | Included (n=42) | Dropout vs Included | Dropout (n=41) | Included (n=24) | Dropout vs Included |
| Sex (M/F) | 19/35 | 22/20 | $\chi^2(1) = 2.2$, $P$ = .14 | 32/9 | 19/5 | $\chi^2(1) = 0$, $P$ = 1 |
| Age | 23.24 (4.29) | 23.93 (3.39) | $T(94)$ = -0.852, $P$ = .4 | 22.12 (4.57) | 21.88 (4.00) | $T(63)$ = 0.22, $P$ = .83 |
| Ethnicity (B/EA/H/O/W) | 38/0/3/1/12 | 23/2/0/2/15 | $\chi^2(4) = 7.98$, $P$ = .092 | 18/2/0/4/17 | 17/0/0/1/6 | $\chi^2(3) = 4.98$, $P$ = .17 |
| Handedness (R/L) | 51/3 | 35/7 | $\chi^2(1) = 2.05$, $P$ = .15 | 39/2 | 21/3 | $\chi^2(1) = 0.398$, $P$ = .53 |
| Smoker (Yes/No) | 1/53 | 3/39 | $\chi^2(1) = 0.596$, $P$ = .44 | 16/25 | 5/19 | $\chi^2(1) = 1.53$, $P$ = .22 |
| Cannabis (Yes/No) | 4/50 | 3/39 | $\chi^2(1) = 0$, $P$ = 1 | 13/28 | 4/20 | $\chi^2(1) = 1.08$, $P$ = .3 |
| Duration of Illness (weeks) | | | | 69.33 (69.33)† | 65.00 (46.58)† | $T(63)$ = 0.319, $P$ = .75 |
| CPZ (mg) | | | | 262.55 (292.04) | 253.33 (234.29) | $T(63)$ = 0.132, $P$ = .9 |
| SAPS | | | | 3.83 (3.40) | 4.48 (4.36) | $T(61)$ = -0.661, $P$ = .51 |

| | Healthy Control | | | Early Psychosis | | |
|---|---|---|---|---|---|---|
| | Dropout (n=54) | Included (n=42) | Dropout vs Included | Dropout (n=41) | Included (n=24) | Dropout vs Included |
| SANS | | | | 9.10 (5.70) | 8.43 (4.04) | $T(61) =$ 0.492, $P$ = .62 |

† Median (IQR)
B=Black/African; EA=East Asian; O=Other/Unknown; W=White/European; CPZ=chlorpromazine equivalent dose; SAPS=Scale for assessment of positive symptoms; SANS=Score for assessment of negative symptoms

Table S4: TOPSY dropout demographics.

| | Healthy Control | | | First Episode Psychosis | | |
|---|---|---|---|---|---|---|
| | Dropout (n=24) | Included (n=15) | Dropout vs Included | Dropout (n=52) | Included (n=19) | Dropout vs Included |
| Sex (M/F) | 16/8 | 10/5 | $\chi^2(1) = 0$, $P = 1$ | 41/10 | 16/3 | $\chi^2(1) = 0.00039$, $P$ = .98 |
| Age | 21.83 (3.80) | 21.73 (2.99) | $T(37) = 0.0865$, $P$ = .93 | 22.74 (4.07) | 22.53 (5.42) | $T(67) = 0.177$, $P$ = .86 |
| Handedness (R/L/A) | 22/0/2 | 14/0/1 | $\chi^2(1) = 0$, $P = 1$ | 41/1/10 | 18/0/1 | $\chi^2(2) = 2.54$, $P$ = .28 |
| Education | 14.04 (2.23) | 14.27 (2.02) | $T(36) = -0.313$, $P$ = .76 | 12.62 (2.06) | 12.95 (1.18) | $T(67) = -0.652$, $P$ = .52 |
| SES | 3.09 (1.20) | 3.20 (1.57) | $T(36) = -0.251$, $P$ = .8 | 3.49 (1.60) | 3.89 (1.08) | $T(57) = -0.97$, $P$ = .34 |
| CAST | 7.17 (3.28) | 7.00 (3.87) | $T(37) = 0.144$, $P$ = .89 | 12.44 (5.95) | 12.12 (6.90) | $T(59) = 0.177$, $P$ = .86 |
| AUDIT-C | 3.38 (2.24) | 2.87 (2.26) | $T(37) = 0.686$, $P$ = .5 | 1.60 (2.19) | 3.79 (3.83) | $T(54) = -2.65$, **$P$ = .01** |

| | Healthy Control | | | First Episode Psychosis | | |
|---|---|---|---|---|---|---|
| | Dropout (n=24) | Included (n=15) | Dropout vs Included | Dropout (n=52) | Included (n=19) | Dropout vs Included |
| Smoker (yes/no) | 0/24 | 1/14 | $\chi^2(1) = 0.0577$, $P$ = .81 | 12/40 | 7/12 | $\chi^2(1) = 0.735$, $P$ = .39 |
| Cannabis (yes/no) | 7/17 | 5/10 | $\chi^2(1) = 0$, $P = 1$ | 31/15 | 12/7 | $\chi^2(1) = 0.00159$, $P$ = .97 |
| SOFAS | 82.52 (3.14) | 81.08 (6.24) | $T(32) = 0.9$, $P$ = .37 | 39.71 (12.20) | 42.00 (13.37) | $T(69) = -0.682$, $P$ = .5 |
| Duration of Illness (weeks) | | | | 139.00 (238.00)[†] | 104.00 (132.50)[†] | $T(53) = 1.02$, $P$ = .31 |
| Antipsychotics (DDD-days) | | | | 0.20 (0.66)[†] | 0.00 (0.33)[†] | $T(66) = 0.551$, $P$ = .58 |
| PANSS-8 Total | | | | 25.71 (7.70) | 24.68 (5.53) | $T(62) = 0.526$, $P$ = .6 |
| PANSS-8 Positive | | | | 12.18 (3.14) | 12.53 (2.82) | $T(62) = -0.417$, $P$ = .68 |
| PANSS-8 Negative | | | | 8.00 (4.66) | 6.26 (3.31) | $T(63) = 1.47$, $P$ = .15 |
| PANSS-8 General | | | | 5.57 (2.24) | 5.89 (2.54) | $T(63) = -0.519$, $P$ = .61 |

[†] Median (IQR)
B=Black/African; C=Caribbean/North American Black; EA=East Asian; W=White/European; DDD-days=Defined daily dose × Days; CDS=Calgary Depression Scale; CAST=Cannabis Abuse Screening Test; PANSS=Positive and Negative Symptom Scale; SES=Socioeconomic status; AUDIT-C=Alcohol Use Disorders Identification Test; SOFAS=Social and Occupational Functioning Assessment Scale

Table S5: JHSZC age-matched demographics (age < 24yr).

| | Healthy Control (n=19) | | | Early Psychosis (n=21) | | |
|---|---|---|---|---|---|---|
| | Baseline (n=19) | 1yr (n=13) | 2yr (n=4) | Baseline (n=17) | 1yr (n=14) | 2yr (n=7) |
| Sex (M/F) | 9/10 | 5/8 | 1/3 | 14/3 | 11/3 | 6/1 |
| Age | 21.05 (1.58) | 21.85 (1.34) | 21.00 (0.82) | 19.88 (2.67) | 20.36 (2.41) | 20.14 (2.48) |
| Ethnicity (B/EA/O/W) | 11/1/1/6 | 8/0/1/4 | 2/1/1/0 | 13/0/0/4 | 13/0/0/1 | 6/0/0/1 |
| Handedness (R/L) | 17/2 | 11/2 | 4/0 | 15/2 | 12/2 | 7/0 |
| Smoker (Yes/No) | 0/19 | 0/13 | 0/4 | 4/13 | **6/8** | 2/5 |
| Cannabis (Yes/No) | 1/18 | 1/12 | 0/4 | 3/14 | 3/11 | 0/7 |
| Duration of Illness (weeks) | | | | 56.33 (73.67)$^{\dagger}$ | 106.17 (82.33)$^{\dagger}$ | 160.33 (36.83)$^{\dagger}$ |
| CPZ (mg) | | | | 274.71 (251.69) | 334.29 (280.37) | 342.86 (265.25) |
| SAPS | | | | 4.25 (4.78) | 4.67 (2.87) | 3.86 (5.43) |
| SANS | | | | 8.31 (4.57) | 6.33 (3.87) | 5.86 (6.20) |

$^{\dagger}$ Median (IQR)
B=Black/African; EA=East Asian; O=Other/Unknown; W=White/European;
CPZ=chlorpromazine equivalent dose; SAPS=Scale for assessment of positive symptoms;
SANS=Score for assessment of negative symptoms

Table S6: JHSZC age-matched demographics statistics (age $<$ 24yr).

| | Baseline | 1yr | 2yr |
|---|---|---|---|
| Sex (M/F) | $\chi^2(1) = 3.36$, $P$ = .067 | $\chi^2(1) = 2.98$, $P$ = .084 | $\chi^2(1) = 1.86$, $P$ = .17 |
| Age | $T(34) = 1.62$, $P$ = .11 | $T(25) = 1.96$, $P$ = .061 | $T(9) = 0.658$, $P$ = .53 |
| Ethnicity (B/EA/O/W) | $\chi^2(3) = 2.46$, $P$ = .48 | $\chi^2(2) = 3.96$, $P$ = .14 | $\chi^2(3) = 4.52$, $P$ = .21 |
| Handedness (R/L) | $\chi^2(1) = 0$, $P = 1$ | $\chi^2(1) = 0$, $P = 1$ | $\chi^2(0) = 0$, $P = 1$ |
| Smoker (Yes/No) | $\chi^2(1) = 2.93$, $P$ = .087 | $\chi^2(1) = 4.9$, ***P* = .027** | $\chi^2(1) = 0.136$, $P$ = .71 |
| Cannabis (Yes/No) | $\chi^2(1) = 0.421$, $P$ = .52 | $\chi^2(1) = 0.213$, $P$ = .64 | $\chi^2(0) = 0$, $P = 1$ |

B=Black/African; EA=East Asian; O=Other/Unknown; W=White/European
