## Supplementary Methods for "Stable White Matter Structure in the First Three Years after Psychosis Onset"

### 1 Data

#### 1.1 Johns Hopkins Schizophrenia Center (JHSZC)

This study was approved by the Johns Hopkins Medicine Institutional Review Board. All study participants provided written informed consent. 101 patients and 96 controls were recruited from within Johns Hopkins Hospital and the surrounding region. Inclusion criteria were: 1) between 13 and 35 years old; 2) no history of traumatic brain injury, cancer, abnormal bleeding, viral infection, neurologic disorder, or intellectual disability; 3) no drug or alcohol abuse (not including cannabis or synthetic cannabinoid receptor agonists) in the past three years; 4) no illicit drug use in the past two months. Additional inclusion criteria for patients were: at baseline, within 24 months of the onset of psychotic manifestations as assessed by study team psychiatrists using the Structured Clinical Interview for DSM-IV (SCID) and collateral information from available medical records. All but six patients were already medicated at their first (baseline) visit. The six medication-naive patients started medication during the follow-up period. The study psychiatrists did not make treatment decisions regarding medications.

Five patients were initially diagnosed as substance-induced psychotic disorder because of the use of cannabis or synthetic cannabinoid receptor agonists. However, two of these were later diagnosed as bipolar disorder with psychotic features, and one with schizophrenia. After the enrollment (first visit), controls and patients were followed up to 4 years. From this initial dataset, two final exclusions were made: first, subjects with only one usable session were excluded, and second, only patients with a diagnosis of schizophrenia, schizoaffective disorder, schizophreniform disorder, or psychosis not otherwise specified. Subjects with bipolar disorder with psychotic features (n = 22), major depressive disorder with psychotic features (n = 4), substance induced psychotic disorder (n = 3), and psychosis not otherwise specified (n=3) were excluded.

The Scale for the Assessment of Negative Symptoms and the Scale for the Assessment of Positive Symptoms was used to evaluate the presence and severity of negative and positive symptoms respectively (1).

Follow-up assessments and imaging for both healthy controls (HCs) and patients were obtained at one, two, and three years from baseline, for all subjects available for reassessment.

Data was acquired with a head-only, neuro-optimized 3T MRI (Philips Intera (v3.2.2)) using T1-weighted (T1w) and diffusion magnetic resonance imaging (dMRI) imaging protocols. T1w data was collected using an MPRAGE sequence at 1 mm isotropic resolution, echo time = 3.7 ms, repetition time = 8.1 s, field of view = 224x180x165 mm, number of slices = 165, flip angle=8. Two replicate diffusion datasets were acquired per subject per session with an echo planar imaging (EPI) sequence at 0.74x0.74x2.2 mm resolution, echo time = 75 ms, repetition time = 7.4 s, field of view = 212.53 mm, number of slices = 70, MB acceleration factor = 2, flip angle = 90. 32 directions were acquired in the AP direction at b=700, along with 1 b=0 images.

The diffusion data from this dataset have not been presented in any previous report.

#### 1.2 Tracking Outcomes in Psychosis (TOPSY)

First episode psychosis (FEP) patients and HCs were recruited from an established cohort enrolled in the Prevention and Early Intervention Program for Psychoses (PEPP) in London, Ontario. This is a high fidelity, early intervention programme that uses an intense case management model, receiving all incident cases of psychosis in the defined catchment area, with first contact made within 48 hours of referral. Inclusion criteria for FEP patients was as follows: individuals experiencing their first psychotic episode, with no more than 14 days of cumulative lifetime antipsychotic exposure, ability to provide informed consent, no major head injuries (leading to a significant period of unconsciousness or seizures), no known neurological disorders, and no concurrent substance use disorder. Participants were not explicitly instructed to abstain from substances, and patients on non-antipsychotic prescription medication were not excluded. All participants provided written, informed consent prior to participation as per approval provided by the Western University Health Sciences Research Ethics Board, London, Ontario. Patient consensus diagnosis was established using the best estimate procedure described by Leckman *et al.* (2) and confirmed after 6 months of treatment. The diagnosis of schizophrenia was based on the DSM-5 criteria.

HCs were recruited through posters and word-of-mouth advertising. They had no personal history of mental illness, no current use of medications, and no first-degree relatives with a history of psychotic disorders. HCs were group matched to the FEP cohort for age and sex. Like their FEP counterparts, those with a history of substance use disorders in the past twelve months, significant head injury, or neurological disorders were excluded.

Four FEP subjects that did not receive a schizophrenia spectrum disorder diagnoses (two with major depressive disorder, two with bipolar disorder) were excluded from analysis.

Follow-up assessments and imaging for both HCs and patients were obtained at six months, one year, two years, and three years, for all subjects available for reassessment. Only one follow-up was ever attained for any of our HCs, typically at the six month or one year time point. Thus, the analysis was conducted using data from the baseline and first follow-up assessment for all subjects.

Data was acquired with a head-only, neuro-optimized 7T MRI (Siemens MAGNETOM Plus, Erlangen, Germany) using dMRI and T1w imaging protocols. T1w data was collected using an MP2RAGE sequence (3) at 0.75 mm isotropic resolution, echo time = 2.83 ms, repetition time = 6 s, field of view = 240x240 mm, number of slices = 208. The T1w image was reconstructed using the robust algorithm introduced by O'Brien et al (4). Diffusion data was acquired with an EPI sequence at 2mm isotropic resolution, echo time = 50.2 ms, repetition time = 5.1 s, field of view = 208 mm, number of slices = 72, MB acceleration factor = 2, flip angle = 90. 64 directions were acquired in both the AP and PA directions at b=1000, along with 2 b=0 images. Gradient nonlinearity correction was applied to all acquisitions using in-house software.

The baseline diffusion data from this dataset was previously analyzed (5). In this report, baseline data is compared to previously unreported follow-up data.

### 2 Preprocessing

Anatomical and diffusion preprocessing were performed as previously reported (5), briefly summarised below.

#### 2.1 Anatomical data

Segmentation of anatomical images and construction of the cortical surface mesh was performed using *FastSurfer* (6–8). These outputs were further processed with *ciftify* (9), an implementation of the Human Connnectome Project minimal preprocessing workflow (10). Images were registered to the *MNI152NLin6Asym* (11) template space, and meshes were registered to the *fsLR-32k* template space (12).

#### 2.2 Diffusion Data

Diffusion data was preprocessed using *snakedwi* (13), a preprocessing pipeline based on *snakebids* (14) and *snakemake* (15). Briefly, Gibbs ringing artefacts were removed with *mrdegibbs* from *MRtrix3* (16,17); eddy currents and motion were corrected using *eddy* from *FSL* (18); susceptibility-induced distortions were corrected using *topup* in *FSL*, using the AP-PA pairs of images (19,20). The T1w image was skull stripped with *SynthStrip* (21); bias field correction was applied with *N4ITK* from *ANTS* (22). A T1w proxy image was created from the diffusion image using *SynthSR* (23) and used to accurately register the diffusion image space to the T1w space using a rigid transform calculated with *greedy* (24). Diffusion tensor imaging (DTI) metrics were calculated using *dtifit* from *FSL* with linear regression (25).

Tractography was performed using the *MRtrix3* (17) software suite. Constrained spherical deconvolution was performed using the dhollander algorithm to estimate the response functions for white matter, grey matter and cerebrospinal fluid (CSF) (26,27). Single shell 3 tissue constrained spherical deconvolution (SS3T-constrained spherical deconvolution), as implemented in *MRtrix3Tissue* (https://3tissue.github.io/), was performed to obtain white matter-like fibre orientation distributions (FODs) as well as grey matter-like and CSF-like compartments in all voxels (28). *mtnormalise* was used to correct for residual intensity inhomogeneities (29,30). Each FOD then models the directionally-specific diffusion signal attributable to white matter within the voxel, increasing the robustness of future processing to crossing tracts (31). Tractography was performed using the iFOD2 algorithm (32) and anatomically constrained tractography (ACT) (33), with 10,000,000 streamlines (34), an FOD amplitude cut-off of 0.06, a minimum streamline length of 4mm, and a maximum streamline length of 250mm. The anatomical segmentation used for ACT was obtained using *SynthSeg* (35).

Subject anatomical scans were parcellated using the Brainnetome atlas (36). These parcellations were used to derive weighted connectivity matrices based on the tractography data. Average fractional anisotropy (FA), mean diffusivity, radial diffusivity, axial diffusivity, and the log-transformed spherical-deconvolution informed filtering of tractograms-weighted streamline count were used as weights.

#### 2.3 JHSZC processing

JHSZC data were preprocessed as with TOPSY, with the following changes:

The replicate diffusion acquisitions were concatenated before processing.

Because we did not have a reverse phase-encoding scan, susceptibility distortion correction was performed as follows. First, the T1w image was rigid transformed to the diffusion b0 image. A T1w proxy image was created from the diffusion b0 images using *SynthSR* (23) and registered to the aligned T1w image with *antsRegistration* (37). The inverse warp field from this registration was used to unwarp the distorted diffusion images.

Slice to volume correction in *eddy* was used to correct intravolume motion correction. Following correction, all diffusion volumes were visually inspected for uncorrected motion artefacts (e.g. “zebra-striping”).

Volumes with (qualitatively) severe artefacts were manually excluded. Because of the replicate data sets, this exclusion did not cause bias in the sampling directions.

Because of the low angular resolution of the JHSZC data, we did not perform tractography. Thus, we did not need to transform the diffusion data into the T1w space, so the DTI tensor was fit in the original acquisition space.

No further processing was performed on the anatomical scans. They were not used in subsequent analyses.

### 3 Skeletonization

FA maps were first non-linearly registered to a common template corresponding to the average space of all FA images. This template was computed with an in-house implementation of the iterative algorithm described by Avants *et al.* (38), using *greedy* (24) to compute each transformation. Neurite orientation dispersion and density imaging derived parameter maps (neurite density index (NDI), orientation dispersion index (ODI), $\log \nu_{iso}$) were transformed into this space. *FSL* (20) was then used to skeletonize the maps.

### 4 Parcellations

Global surface averages were computed by averaging across all vertices for each parameter map, per hemisphere. The Desikan-Killiany atlas computed by *fastsurfer* was used to parcellate the surface data. Hemispheres were kept separate throughout the analysis.

A composite, hierarchical atlas was constructed using the Johns Hopkins University (JHU) atlas distributed with *FSL* (39) and the Talairach lobe segmentation (40). Both atlases were first intersected with the skeletonized image, and the lobe segmentation was used to define a peripheral atlas by subtracting the JHU atlas. Mathematically:

$$A^{JHU} = JHU \cap M^{skeleton}$$

$$A^{Lobe} = Lobe \cap M^{skeleton}$$

$$A^{Periph}_{ijk} = \begin{cases} A^{Lobe}_{ijk} & A^{JHU}_{ijk} = 0 \\ 0 & A^{JHU}_{ijk} \neq 0 \end{cases}$$

where $A_{ijk}$ is the $ijk$th voxel of the indicated atlas. Homologous ROIs across the two hemispheres were merged.

Four hierarchical levels were defined for statistical purposes:

1. The global mean (1 ROI)
2. Mean of all voxels in the peripheral atlas and of all voxels in the JHU atlas (2 ROIs)
3. Peripheral atlas and JHU atlas merged into 4 tract groupings (11 ROIs)
4. JHU atlas (23 ROIs)

### 5 References

1. Andreasen NC, Flaum M, Arndt S, Alliger R, Swayze VW (1991): Positive and Negative Symptoms: Assessment and Validity. In: Marneros A, Andreasen NC, Tsuang MT, editors. *Negative Versus Positive Schizophrenia*. Berlin, Heidelberg: Springer, pp 28–51.
2. Leckman JF, Sholomskas D, Thompson WD, Belanger A, Weissman MM (1982): Best estimate of lifetime psychiatric diagnosis: a methodological study. *Archives of General Psychiatry* 39: 879–883.
3. Marques JP, Kober T, Krueger G, Zwaag W van der, Van de Moortele P-F, Gruetter R (2010): MP2RAGE, a self bias-field corrected sequence for improved segmentation and T1-mapping at high field. *NeuroImage* 49: 1271–1281.
4. O'Brien KR, Kober T, Hagmann P, Maeder P, Marques J, Lazeyras F, *et al.* (2014): Robust T1-Weighted Structural Brain Imaging and Morphometry at 7T Using MP2RAGE. *PLOS ONE* 9: e99676.
5. Van Dyken PC, MacKinley M, Khan AR, Palaniyappan L (2024): Cortical Network Disruption Is Minimal in Early Stages of Psychosis. *Schizophrenia Bulletin Open* 5: sgae010.
6. Faber J, Kügler D, Bahrami E, Heinz L-S, Timmann D, Ernst TM, *et al.* (2022): CerebNet: A Fast and Reliable Deep-Learning Pipeline for Detailed Cerebellum Sub-Segmentation. *NeuroImage* 264: 119703.
7. Henschel L, Conjeti S, Estrada S, Diers K, Fischl B, Reuter M (2020): FastSurfer - A Fast and Accurate Deep Learning Based Neuroimaging Pipeline. *NeuroImage* 219: 117012.
8. Henschel L, Kügler D, Reuter M (2022): FastSurferVINN: Building Resolution-Independence into Deep Learning Segmentation methods—A Solution for HighRes Brain MRI. *NeuroImage* 251: 118933.
9. Dickie EW, Smith DE, Mathu-M, lgrennan, Jeyachandra J, delaneyjohnston, Gorgolewski C (2019, August 16): Edickie/ciftify: Fix to ciftify_meants and new ciftify_dlabel_to_vol script. Zenodo. https://doi.org/10.5281/zenodo.3369937
10. Glasser MF, Sotiropoulos SN, Wilson JA, Coalson TS, Fischl B, Andersson JL, *et al.* (2013): The minimal preprocessing pipelines for the Human Connectome Project. *NeuroImage* 80: 105–124.
11. Evans AC, Janke AL, Collins DL, Baillet S (2012): Brain Templates and Atlases. *NeuroImage* 62: 911–922.
12. Van Essen DC, Glasser MF, Dierker DL, Harwell J, Coalson T (2012): Parcellations and Hemispheric Asymmetries of Human Cerebral Cortex Analyzed on Surface-Based Atlases. *Cerebral Cortex* 22: 2241–2262.
13. Khan A, Kai J, Van Dyken P, Karat B (2023, August): Akhanf/snakedwi: 0.2.1, version v0.2.1. Zenodo. https://doi.org/10.5281/zenodo.8237288

14. Khan A, Van Dyken P, Kai J, Kuehn T, Gau R (2023, February 6): Akhanf/snakebids: 0.7.2. Zenodo. https://doi.org/10.5281/zenodo.7613561
15. Mölder F, Jablonski KP, Letcher B, Hall MB, Tomkins-Tinch CH, Sochat V, *et al.* (2021, April 19): Sustainable Data Analysis with Snakemake [no. 10:33]. F1000Research. https://doi.org/10.12688/f1000research.29032.2
16. Kellner E, Dhital B, Kiselev VG, Reisert M (2016): Gibbs-Ringing Artifact Removal Based on Local Subvoxel-Shifts. *Magnetic Resonance in Medicine* 76: 1574–1581.
17. Tournier J-D, Smith R, Raffelt D, Tabbara R, Dhollander T, Pietsch M, *et al.* (2019): MRtrix3: A Fast, Flexible and Open Software Framework for Medical Image Processing and Visualisation. *NeuroImage* 202: 116137.
18. Andersson JLR, Sotiropoulos SN (2016): An integrated approach to correction for off-resonance effects and subject movement in diffusion MR imaging. *NeuroImage* 125: 1063–1078.
19. Andersson JLR, Skare S, Ashburner J (2003): How to correct susceptibility distortions in spin-echo echo-planar images: application to diffusion tensor imaging. *NeuroImage* 20: 870–888.
20. Smith SM, Jenkinson M, Woolrich MW, Beckmann CF, Behrens TEJ, Johansen-Berg H, *et al.* (2004): Advances in Functional and Structural MR Image Analysis and Implementation as FSL. *NeuroImage* 23: S208–S219.
21. Hoopes A, Mora JS, Dalca AV, Fischl B, Hoffmann M (2022): SynthStrip: Skull-stripping for any brain image. *NeuroImage* 260: 119474.
22. Tustison NJ, Avants BB, Cook PA, Zheng Y, Egan A, Yushkevich PA, Gee JC (2010): N4ITK: improved N3 bias correction. *IEEE transactions on medical imaging* 29: 1310–1320.
23. Iglesias JE, Billot B, Balbastre Y, Tabari A, Conklin J, Gilberto González R, *et al.* (2021): Joint super-resolution and synthesis of 1 mm isotropic MP-RAGE volumes from clinical MRI exams with scans of different orientation, resolution and contrast. *NeuroImage* 237: 118206.
24. Yushkevich PA, Pluta J, Wang H, Wisse LEM, Das S, Wolk D (2016): IC-P-174: Fast Automatic Segmentation of Hippocampal Subfields and Medial Temporal Lobe Subregions In 3 Tesla and 7 Tesla T2-Weighted MRI. *Alzheimer's & Dementia* 12: P126–P127.
25. Behrens Tej, Woolrich Mw, Jenkinson M, Johansen-Berg H, Nunes Rg, Clare S, *et al.* (2003): Characterization and Propagation of Uncertainty in Diffusion-Weighted MR Imaging. *Magnetic Resonance in Medicine* 50: 1077–1088.
26. Dhollander T, Raffelt D, Connelly A (2016): Unsupervised 3-tissue response function estimation from single-shell or multi-shell diffusion MR data without a co-registered T1 image.
27. Dhollander T, Mito R, Raffelt D, Connelly A (2019): Improved white matter response function estimation for 3-tissue constrained spherical deconvolution.

28. Dhollander T, Connelly A (2016): A novel iterative approach to reap the benefits of multi-tissue CSD from just single-shell (+b=0) diffusion MRI data.
29. Raffelt D, Dhollander T, Tournier J-D, Tabbara R, Smith R, Pierre E, Connelly A (2017): Bias Field Correction and Intensity Normalisation for Quantitative Analysis of Apparent Fibre Density.
30. Dhollander T, Tabbara R, Rosnarho-Tornstrand J, Tournier J-D, Raffelt D, Connelly A (2021): Multi-tissue log-domain intensity and inhomogeneity normalisation for quantitative apparent fibre density.
31. Dell'Acqua F, Tournier J-Donald (2019): Modelling white matter with spherical deconvolution: How and why? *Nmr in Biomedicine* 32. https://doi.org/ggjqcc
32. Tournier J-D, Calamante F, Connelly A (2010): Improved probabilistic streamlines tractography by 2nd order integration over fibre orientation distributions. *Proc Intl Soc Mag Reson Med (ISMRM)* 18.
33. Smith RE, Tournier J-D, Calamante F, Connelly A (2012): Anatomically-Constrained Tractography: Improved Diffusion MRI Streamlines Tractography Through Effective Use of Anatomical Information. *NeuroImage* 62: 1924–1938.
34. Newlin NR, Rheault F, Schilling KG, Landman BA (2023): Characterizing Streamline Count Invariant Graph Measures of Structural Connectomes. *Journal of Magnetic Resonance Imaging* 58: 1211–1220.
35. Billot B, Greve DN, Puonti O, Thielscher A, Van Leemput K, Fischl B, *et al.* (2023): SynthSeg: Segmentation of Brain MRI Scans of Any Contrast and Resolution Without Retraining. *Medical Image Analysis* 86: 102789.
36. Fan L, Li H, Zhuo J, Zhang Y, Wang J, Chen L, *et al.* (2016): The Human Brainnetome Atlas: A New Brain Atlas Based on Connectional Architecture. *Cerebral Cortex* 26: 3508–3526.
37. Avants BB, Epstein CL, Grossman M, Gee JC (2008): Symmetric diffeomorphic image registration with cross-correlation: evaluating automated labeling of elderly and neurodegenerative brain. *Medical Image Analysis* 12: 26–41.
38. Avants BB, Yushkevich P, Pluta J, Minkoff D, Korczykowski M, Detre J, Gee JC (2010): The Optimal Template Effect in Hippocampus Studies of Diseased Populations. *NeuroImage* 49: 2457.
39. Mori S, Wakana S, Zijl PCM van, Nagae-Poetscher LM (2005): *MRI Atlas of Human White Matter*. Elsevier.
40. Talairach J, Szikla G (1980): Application of stereotactic concepts to the surgery of epilepsy. *Acta Neurochirurgica Supplementum* 30: 35–54.
